# A Non-Canonical Raf Function Is Required for Dorsal-Ventral Patterning During *Drosophila* Embryogenesis

**DOI:** 10.1101/2021.07.29.454294

**Authors:** Jay B. Lusk, Ellora Hui Zhen Chua, Prameet Kaur, Isabelle Chiao Han Sung, Wen Kin Lim, Vanessa Yuk Man Lam, Nathan Harmston, Nicholas S. Tolwinski

## Abstract

Proper embryonic development requires directional axes to pattern cells into embryonic structures. In *Drosophila*, spatially discrete expression of transcription factors determines the anterior to posterior organization of the early embryo, while the Toll and TGFβ signalling pathways determine the early dorsal to ventral pattern. Embryonic MAPK/ERK signaling contributes to both anterior to posterior patterning in the terminal regions and to dorsal to ventral patterning during oogenesis and embryonic stages. Here we describe a novel loss of function mutation in the Raf kinase gene, which leads to loss of ventral cell fates as seen through the loss of the ventral furrow, the absence of Dorsal/NFκB nuclear localization, the absence of mesoderm determinants Twist and Snail, and the expansion of TGFβ. Gene expression analysis showed cells adopting ectodermal fates much like loss of Toll signaling. Our results combine novel mutants, live imaging, optogenetics and transcriptomics to establish a novel role for Raf, that appears to be independent of the MAPK cascade, in embryonic patterning.

## Introduction

A central question of embryonic development is how fates are assigned to thousands of cells within the three-dimensional structure of an egg or embryo. Along the dorsal-ventral axis, the early *Drosophila* embryonic cell fates are determined by the Toll and Dpp/TGFβ pathways, where cells with active Toll adopt a ventral cell fate, and cells with active TGFβ adopt a dorsal cell fate [1–3]. EGF signaling patterns the neuroectoderm in more lateral cells [4], and the combination of EGF, TGFβ and Toll signals establish the dorsal-ventral (D/V) axis in early embryos [4–7].

The EGF signaling pathway is one of the most studied signal transduction cascades in development and disease. The signaling cascade begins with *Epidermal Growth Factor (EGF) receptor (EGFR* or *Der* in *Drosophila)* a transmembrane receptor tyrosine kinase which, among other functions, activates the small GTPase Ras, initiating the Mitogen Activated Protein Kinase (MAPK) cascade through the activation of the Raf kinase [8, 9]. *EGFR*’s *faint little ball* phenotype, which is characterized by the loss of ventral and terminal structures, highlights the importance of the pathway and its components in assigning cell fates [10–13].

The Raf kinase acts downstream of multiple receptor tyrosine kinases in establishing *Drosophila*’s body axes. For example, Raf’s kinase activity downstream of the *torso* receptor tyrosine kinase (RTK) is essential for patterning anterior and posterior termini in the embryo [14, 15]. In follicle cells, the *Torso-like* ligand is only expressed in polar populations. Thus, during embryogenesis, Torso RTK patterning is localized to terminal regions. During early development of the embryo, the ligand binds to Torso and activates Raf, leading to transcriptional activation of zygotic genes *tailless*, *huckebein*, and repression of *bicoid* target genes, such as *hunchback* and *orthodenticle*. The genes then determine cellular fates at the anterior/posterior termini, giving rise to specialized structures at the terminal regions of the larva [16].

Embryos lacking either maternal *Raf* or *Tor* activities demonstrate significant aberrations in anteroposterior patterning, including loss of hindgut and posterior midgut structures, derivatives of abdominal segments 8 to 10, specific head skeletal structures, and Malphigian tubules [17–19].

Raf also acts downstream of EGFR to determine ventral ectodermal and lateral fates during embryogenesis [20–22]. Without EGFR activity, embryos will display the *faint little ball* phenotype, characterized by the presence of only the dorsal hypoderm in cuticles and a loss of ventral and terminal structures [10–13]. As Raf works downstream of EGFR in developing the ventral ectoderm, a constitutively active Raf can bypass defective *EGFR* and rescue the *faint little ball* phenotype [15].

Raf also plays other roles in development. For instance, Raf acts upstream of *seven-in-abstentia* (*sina*) and downstream of Ras and receptor tyrosine kinase *sevenless* (*sev*) in the Sev pathway to establish R7 photoreceptor cells in the eye [23–25]. Constitutively activated Raf can induce R7 development in the *sevenless* phenotype that lacks R7 in every ommatidium of the eye [24]. After development, EGFR signaling plays a multitude of roles ranging from wound healing to homeostasis in both vertebrates and *Drosophila* [26–28].

Raf is also essential for establishing dorsoventral cell fates during oogenesis. During oogenesis, the asymmetrically anchored oocyte nucleus defines a presumptive dorsal region that contains high levels of Gurken, a transforming growth factor (TGF-α) ligand that activates EGFR in the dorsal oocyte follicular epithelium [29–32]. The activating signal from Gurken is then transduced through the classical MAPK cascade, leading to the transcription of a variety of target genes, such as *rhomboid* [33]. Rhomboid is then apically localized in the dorsal follicle cells on the anterior side of the egg chamber. It is required to induce dorsal cell fates in the oocyte and consequently in the embryo [34]. *Rhomboid* also activates another EGFR ligand, Spi, through an intracellular cleavage mechanism [33]. At the maximum level of EGF signaling, located at the dorsal midline, the EGF repressor *argos* is expressed, with the end result being two stripes of *rhomboid* expression which surround the dorsal midline, thereby generating a complex dorsal-ventral pattern in the follicle cells which will later influence embryonic development [35, 36].

The Gurken activated EGF pathway represses *pipe* in the dorsal region follicle layer of the oocyte [31, 37]. When the egg is fertilized, Pipe in the ventral region of the embryo facilitates a proteolytic cascade that results in the release of Spätzle. Spätzle then activates the Toll pathway receptor and delimits and orients the dorsoventral axis of the embryo [38]. Above all, EGF signaling in the maternal follicle cells is required to repress ventral cell fates and is sufficient to induce dorsal cell fates in the oocyte. More specifically, contacts between the follicle cells and the oocyte during oogenesis can cause limited activation of EGFR in the oocyte, which represses ventral cell fates in the developing oocyte [33, 39, 40]. This initial dorsal-ventral polarity of the oocyte later becomes important in the dorsal-ventral patterning of the fertilized embryo.

Once the Toll receptor has been activated by <u>Spätzle</u>, the *Drosophila* homologue of myeloid differentiation primary response 88 (*dMyD88* or *krapfen (kra))* binds directly to Toll and interacts with the death domain of a downstream protein called Tube [41, 42]. Tube, the orthologue of mammalian Interleukin-1 receptor associated kinase 4 (IRAK-4), is a short protein containing an N-terminal death-domain, and a C-terminal Tube repeat domain that participates in protein-protein interactions [43]. Unlike mammalian IRAKs, Tube does not have a kinase domain and relies instead on Pelle kinase to transduce the Toll signal. The Tube/MyD88 complex binds to Pelle, Pelle is autophosphorylated, which is required for Toll signal transduction [44, 45]. IκB-α (*Drosophila* Cactus), in the absence of Toll signaling, binds to RelA/NF-κB (*Drosophila* Dorsal) sequestering NF-κB in the cytoplasm, leading to NF-κB degradation [46]. Phosphorylation of IκB-α leads to its degradation, which releases NF-κB, allowing NF-κB to translocate to the nucleus to initiate transcription [47]. In mammals, the IκB kinase (IKK) complex is composed of two kinases (IKKα and IKKβ) and a regulatory NEMO subunit [48]. The canonical (pro-inflammatory response) pathway is activated by phosphorylation of IKKβ, and the non-canonical pathway (associated with lymphoid cell proliferation) is activated by phosphorylation of IKKα, which itself is activated by a distinct upstream kinase called the NF-κB-inducing kinase (NIK), which is a member of the Map Kinase Kinase Kinase family (MAP3K14) [49, 50]. The *Drosophila* Toll pathway uses a different IKK complex [51], where the kinase Pelle can fulfil the functional role of IKKβ [52].

Activation of Toll followed by the phosphorylation and degradation of Cactus/IκB [38, 53] frees Dorsal/NFκB to enter the nucleus and begin transcription [54–57]. Once inside the nucleus, Dorsal activates or represses a wide variety of genes, including key developmental regulators *decapentaplegic, zerknüllt, twist,* and *snail,* which pattern much of the rest of the embryo [7, 58]. Although the centrality of Toll signaling for D/V patterning has long been established, not much is known about signaling cross talk between the Toll pathway and the EGFR/Ras/Raf pathway in early embryogenesis. The canonical functions of Raf require the MAPK cascade, but there are some examples where Raf functions independently [59–62]. Here, we turn to examining the role Raf plays in early patterning events that do not coincide with previously defined domains of MAPK activity by analyzing new mutants.

## Results

### A Dorsalizing *Raf* Mutation

To identify genes involved in D/V patterning, we screened a new library of molecularly defined mutations for defects in axis formation [63]. ∼2000 X-chromosome mutants were independently crossed to generate maternally and zygotically mutant embryos and screened for patterning defects by cuticle phenotypes [64–66]. We discovered a strong dorsalizing mutation (926, Fig. 1D) that was mapped through complementation analysis to a chromosomal region containing the *Raf* gene. Molecular analysis showed that the *Raf^926^* DNA sequence contained a deletion of 17 nucleotides beginning at the 479th nucleotide of the protein-coding region. This lesion created a frame shift in the coding sequence, leading to a premature stop codon before the second and third conserved domains, in addition to deleting the majority of the first conserved domain (Fig.1A). This deletion was predicted to be a strong loss of function allele as the stop codon eliminates most of the Ras-binding domain (RBD) in conserved region 1 (CR1), the negative regulatory domain (NRD) in CR2 and the protein kinase domain in CR3. We were unable to detect mRNA or to make cDNA, likely as a result of nonsense mediated decay [67]. The stop codon is in a similar location to the strong, amorphic Class 1 alleles of *Raf* reported [18]. To confirm that the allele was a loss of function allele and not a dominant negative allele, we cloned genomic DNA from *Raf^926^* into an overexpression construct tagged with mCherry. Overexpression did not show a phenotype in embryonic patterning and development proceeded normally (Combined Videos 1, Video 1). As previous analyses of Raf mutants did not focus on the ventral most cell fates, we proceeded to test more Raf mutants such as the point mutants *Raf^A^* and *Raf^B^*, but these did not produce embryos as embryogenesis was disrupted [63, 68]. To overcome this limitation, we turned to a CRISPR/Cas9 approach where maternally expressed Cas9 drives gRNA directed mutations in the *Raf* gene [69]. We observed dorsalized embryos as shown by the presence of dorsal hairs as well as other patterning phenotypes, which likely depend on the timing of the induced genetic lesion (S. Fig. 1).

**Figure 1.**
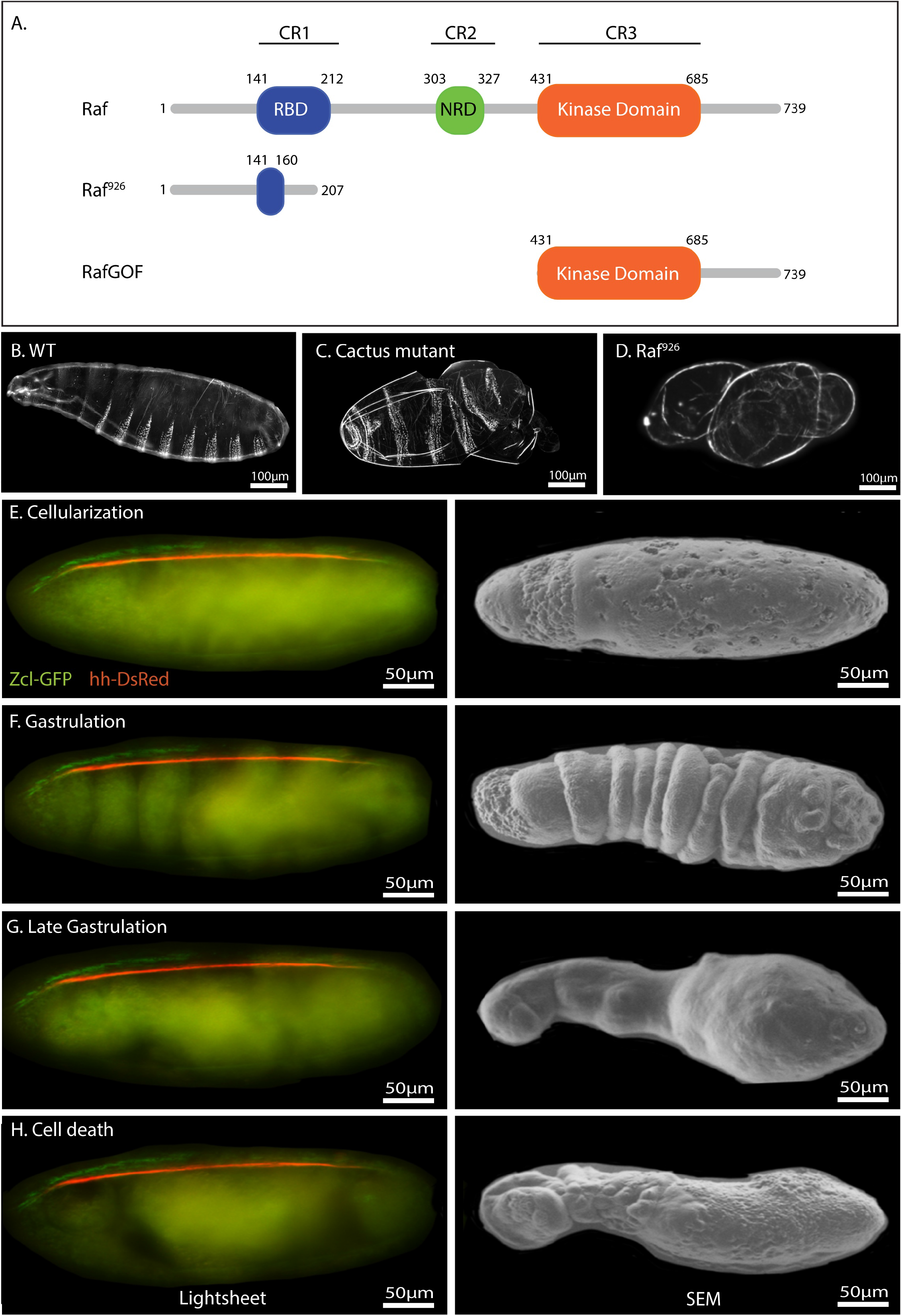
Schematic illustration, cuticular phenotype and developmental progression of *Raf^926^* mutant embryo. **(A)** Domain organization of *Drosophila* Raf, Raf^926^ and Raf^GOF^. Raf protein contains the Ras-binding domain (RBD) in conserved region 1 (CR1), negative regulatory domain (NRD) in CR2 and protein kinase domain in CR3, with a total length of 739 amino acids. Raf^926^ comprised of an altered RBD with 207 amino acids in total due to a deletion of 17 nucleotides which resulted in a frameshift and premature stop codon. Raf^GOF^ contains the kinase domain only, with total length of 309 amino acids. Dark field micrographs of the cuticles of wild type embryo showing normal distribution of ventral denticles and dorsal hair **(B)**, a *cactus^1^* mutant embryo showing strongly ventralized phenotype with ventral denticles expression among the dorsal hair **(C)** and *Raf^926^* mutant embryo showing a strongly dorsalized phenotype with elongated, tube-like, twisted body entirely covered by dorsal hair **(D)**. **(E – H)** Still images of developmental stages of *Raf^926^* embryo from lightsheet (left panel) and scanning electron microscope (SEM) (right panel). *Raf^926^* embryos develop normally up to cellularization stage. Defective gastrulation is characterized by frequent twisting, elongation of twisted segments, fusions of segments into three main sacs, followed by tissue death. Embryo was visualised with Utrophin-GFP and Histone-RFP, with anterior to the left. Stills were extracted from Video 3.

The prototypical dorsalizing mutations such as *dorsal* and *Toll* show an elongated, tube-like embryo that twists within the eggshell and lacks all ventral features such as denticles [2, 70]. *Raf^926^* showed a similar highly-dorsalized phenotype where the embryos appear to twist around themselves and fail to develop ventral structures such as denticles (Fig. 1D). As EGF is involved in oocyte patterning, we inspected ovarioles and egg morphology to determine if any abnormalities were present [32, 71]. We observed that ovarioles were unaffected and oocytes developed standard shape and polarity including normal dorsal appendages. We concluded that the embryonic phenotype was not due to defects in oogenesis, but rather was due to altered signaling in embryogenesis.

To observe early developmental stages of these mutants, we used a live imaging approach through lightsheet microscopy [72, 73]. *Raf^926^* embryos appear morphologically normal through the cellularization stage (Fig. 1E) but begin to show abnormalities during the first process of gastrulation (Fig. 1F), where no ventral furrow is formed. Without a ventral furrow, cells cannot enter the interior of the embryo to form the mesoderm, and the cell movements of gastrulation in dorsalized embryos become unpredictable (Fig. 1G, H, Combined Video 1, Videos 2 and 3 compare to wild-type Video 4). Nevertheless, we looked at the various stages from live imaging and compared these to fixed embryos in scanning electron microscopy to show the various stages of tissue movements (Fig. 1E - H, Video 3). These showed a twisting and elongation of the ectodermal tissue (Fig.1E-H) without the ventral denticle structures that would be seen in a ventralizing mutation such as *cactus* (Fig. 1C).

The primary ventral feature in early stages of gastrulation is the ventral furrow. It is one of the first morphogenetic events of gastrulation and leads to the internalization of mesodermal precursors. Gastrulation starts immediately after cellularization and results in a furrow along the ventral midline [74]. Apical constriction occurs in a 12 cell-wide region surrounding the ventral midline, and these midventral cells then drag three rows of cells on each side towards the ventral midline. As the midventral cells continue to contract, neighboring lateral cells are continuously dragged into the furrow [75, 76]. To determine whether *Raf* loss-of-function could cause disruptions to ventral development during pre- to early gastrulation, we looked at brightfield Videos to follow development of the ventral side of *Raf^926^* embryos starting from before stage 5 to stages 6 or 7. Both wildtype and *Raf^926^* appeared similar during cellularization (Combined Video 2, Compare Video 5 to Video 6, Still Images in S. Fig. 2), but after cellularization, the wildtype embryo formed a clear ventral furrow whereas *Raf^926^* did not (Video 6, Still Images in S. Fig. 2).

Given the early role of EGFR signaling in terminal structure patterning (downstream of *torso* and *torsolike* [19, 77]), we investigated whether disrupted anterior to posterior (A/P) patterning could explain the loss of the ventral furrow. *Raf* mutants show a loss of terminal A/P structures as well as D/V patterning defects [17, 18, 78]. To confirm that the lack of ventral furrow was not due to anteroposterior defects caused by Raf’s other role, we used the triple-mutant *bicoid*, *nanos*, *torso-like* (*bnt*) which lacks anteroposterior patterning [79]. Much like wildtype embryos, *bnt* triple mutants developed a ventral furrow (Video 7, Still Images in S. Fig. 2). The presence of ventral furrow in *bnt* and the absence of ventral furrow in *Raf^926^* suggested a role for Raf in ventral morphogenic events that cannot be accounted for by abnormal anterior/posterior patterning.

### Transcriptional analysis of *Raf^926^* mutants identifies changes in key D/V processes and pathways

As the EGFR signaling pathway has not previously been associated with ventral cell fates and previous studies have not detected phosphorylated ERK in ventral cells [8, 80], we investigated the transcriptional changes occurring in the *Raf^926^* mutants. We collected and sequenced mRNA from *Raf^926^* maternally mutant embryos and compared their transcriptional profiles to wild type (WT) embryos (S. Fig. 3A). As expected, total Raf expression was reduced by more than half (S. Fig. 3B) as expected for embryos maternally deficient for Raf crossed to hemizygous males. We found that 5611 genes were significantly differentially expressed (FDR < 10%, fold change >1.5), with 2593 genes upregulated in *Raf^926^* mutant embryos and 3018 downregulated compared to WT (Fig. 2A, Supplemental Table 1).

**Figure 2:**
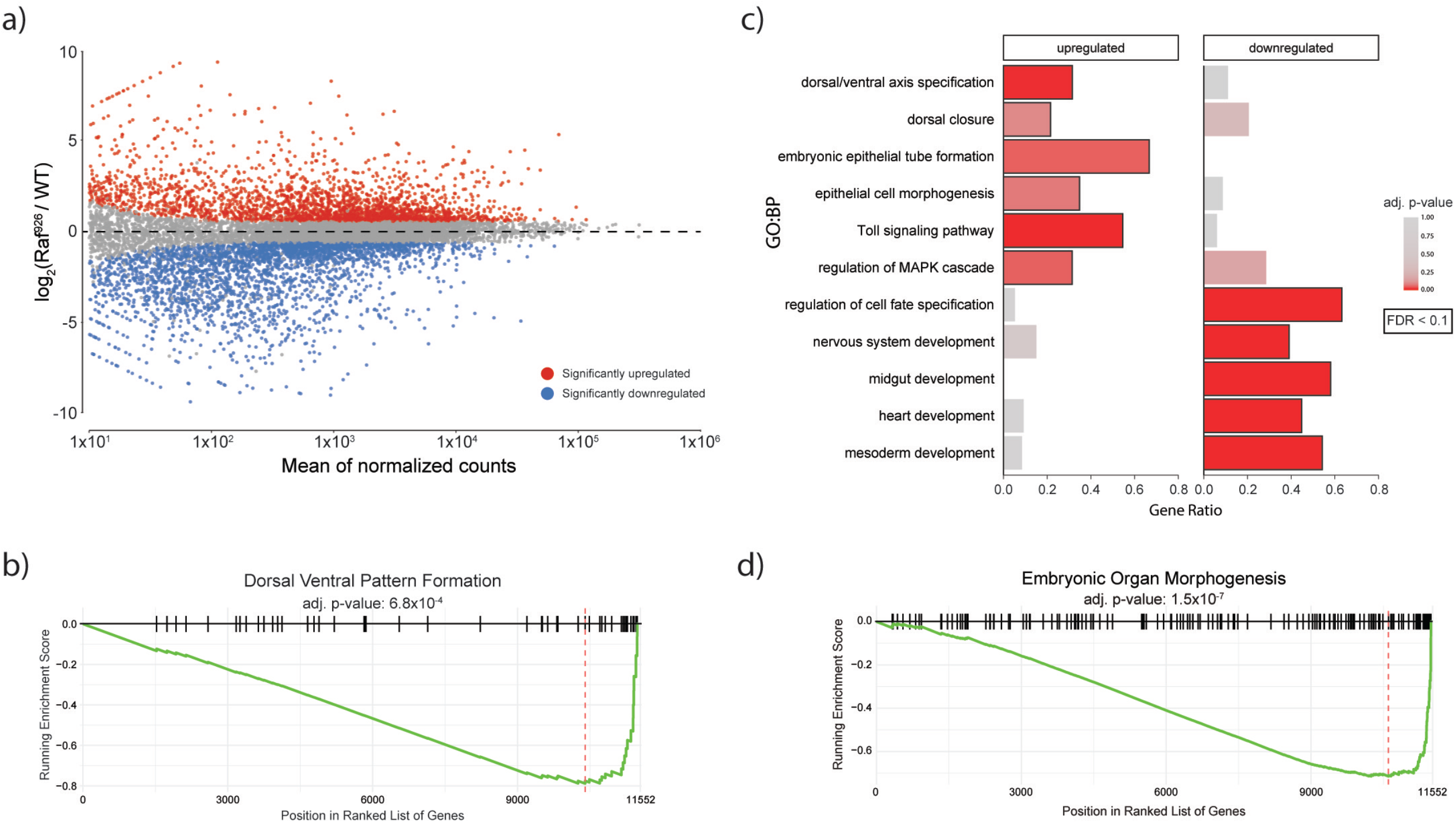
Transcriptional profiling of WT, *Raf^926^* embryos identifies significant differences in the expression of genes involved in Dorsal/Ventral patterning, organogenesis and the Toll pathway. (A) MA-plot shows a large number of genes are significantly differentially expressed between *Raf^926^* mutant and WT embryos. **(B)** GSEA identifies that genes annotated as involved in dorsal-ventral pattern formation are altered in *Raf^926^* mutants. **(C)** GO Biological Process (GO:BP) enrichment analysis identifies that the significant up- and downregulated genes are enriched for distinct processes. **(D)** GSEA identifies that genes are involved in embryonic organ morphogenesis and perturbed in *Raf^926^* mutants.

The expression of genes involved in dorsal-ventral pattern formation was significantly dysregulated in *Raf^926^* mutants (Fig. 2B). Genes that were upregulated in *Raf^926^* mutants were enriched for processes associated with ectoderm, including epithelial cell morphogenesis (FDR < 10%, Fig. 2C). The Toll pathway is required for the formation of ventral cells and TGFβ for dorsal cell fates. Surprisingly, we observed that components of the Toll pathway were upregulated in *Raf^926^* mutants (Fig. 2C, S. Fig. 3C), whereas the set of downregulated genes was enriched for processes related to nervous system and mesoderm development (FDR < 10%, Fig. 2C). In addition, GSEA identified genes involved in embryonic organ morphogenesis as dysregulated in *Raf^926^* embryos (Fig. 2D). These results are consistent with the lack of ventral furrow formation as many internal structures would be missing without the generation of a mesoderm in *Raf^926^* mutants.

### Expression changes in *Raf^926^* mutants are similar to those observed in Toll pathway mutants

To better understand how the transcriptional changes that were observed in the *Raf^926^* mutant reflected differences in germ layer formation, we compared the set of differentially expressed genes with those identified in two distinct datasets investigating germ layer formation during early embryonic development. Stathopoulous *et al.* investigated gene expression changes in three distinct early embryonic regions as defined by levels of Toll signaling [81]. They defined the region with high and medium Toll signalling as mesoderm and neuroectoderm respectively, while the dorsal ectoderm was characterized as the region lacking Toll signalling. Marker genes were subsequently identified for each of these regions. We investigated the effect of *Raf^926^* on the expression of these three cohorts of dorsal/ventral patterning markers. The majority of Dorsal targets (15/20) in the mesoderm were significantly downregulated in *Raf^926^* mutants (Fig. 3B, S. Fig. 3C), including the key mesoderm determinants twist (*twi*) and snail (*sna*) (Fig. 3A). We next looked at Dorsal targets in the neuroectoderm and found that 15/20 targets were significantly downregulated in the *Raf^926^* mutants (Fig. 3B, S. Fig. 4B). In particular, the Dpp pathway regulators *sog* and *brk* (Fig. 3A), as well as the neuroectoderm determining transcription factors intermediate neuroblasts defective (*ind*) and ventral nervous system defective (*vnd*) were downregulated. The third set corresponds to genes that are repressed by Dorsal and as such are expressed on the dorsal side of the embryo which gives rise to the ectoderm. In dorsalized embryos, a condition where no Toll signal is present, these genes increase in expression [81, 82]. We investigated this set of genes and found that 7/13 were significantly upregulated in *Raf^926^* mutants (Fig. 3B, S. Fig. 4A), including key transcription factors involved in dorsal cell fate determination (i.e. pannier (*pnr*) and zerknullt (*zen*), Fig. 3A).

**Figure 3:**
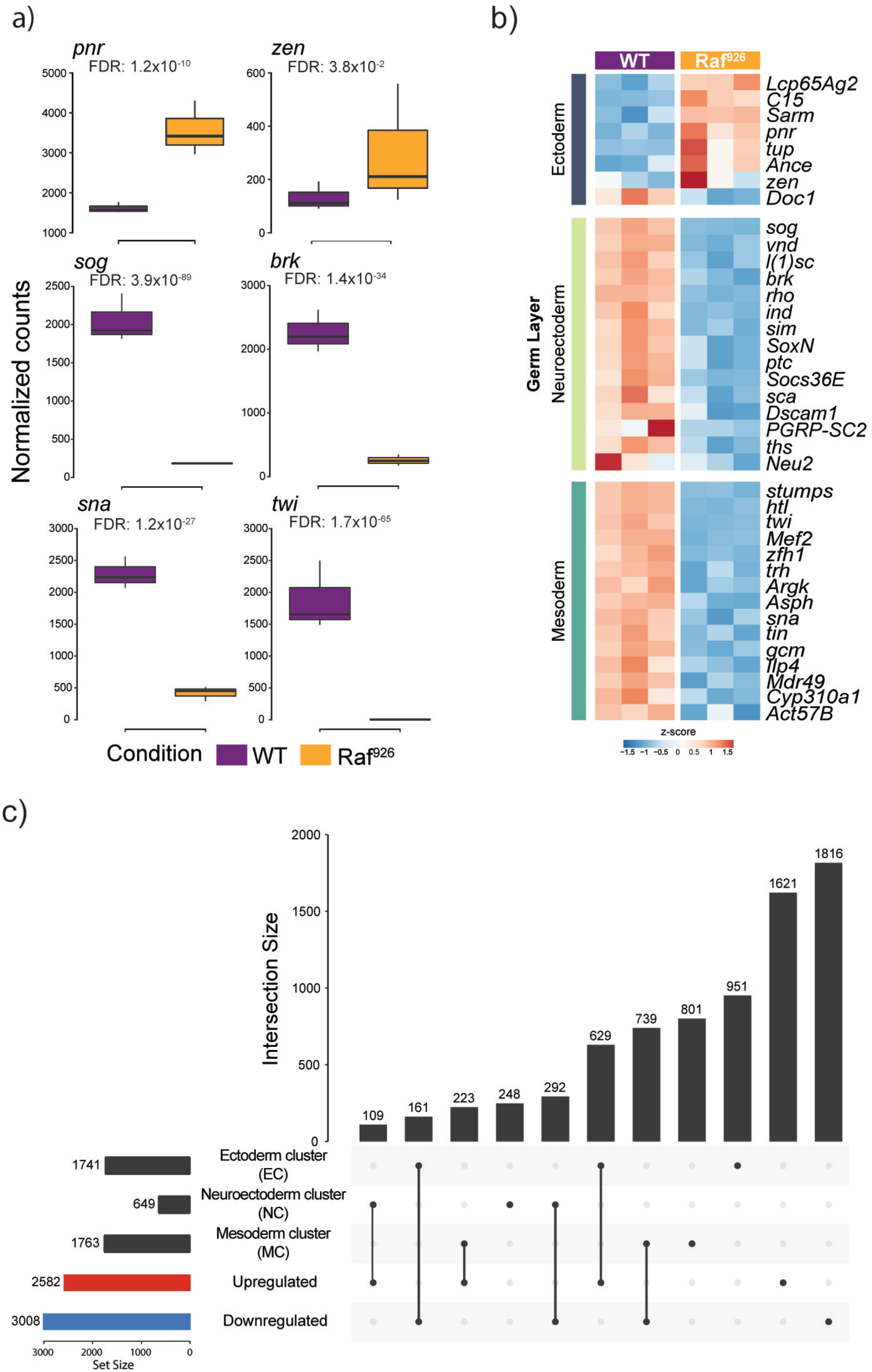
Comparison of expression changes in observed *Raf^926^* mutants with key germ layer marker genes. **(A)** Expression profiles of key germ layer marker genes in WT and *Raf^926^* mutant embryos. **(B)** *Raf^926^* mutants show upregulation of ectoderm marker genes, along with downregulation of neuroectoderm and mesoderm marker genes. **(C)** Upset plot showing the overlap between differentially expressed genes in *Raf^926^* mutant with clusters of gene expression changes observed after generating embryos with only one germ layer after perturbing the Toll pathway (see S. Fig. 3D).

Embryos can be generated which consist of either dorsal ectoderm, neuroectodermal or mesodermal precursor cells using specific mutant lines; gd^7^, Toll^rm9^/^rm10^ and Toll^10b^ respectively. Previously, Koenecke *et al.* characterized the epigenetic and transcriptional landscapes of these mutant embryos [82, 83]. Re- analysis of their RNA-seq dataset identified 4173 significantly differentially expressed genes (FDR < 10%) which clustered into three distinct groups, each associated with distinct germ layers/mutant lines (S. Fig. 4D and E). We compared each of these clusters with the set of significantly differentially expressed genes observed in the *Raf^926^* mutant RNA-seq analysis (Fig. 3C). There was a significant overlap (N=629, p<5x10^-4^) between genes upregulated by *Raf^926^* and those genes located in the cluster of genes specifically upregulated in the dorsal ectoderm (EC). The set of genes downregulated by *Raf^926^* significantly overlapped with genes upregulated in neuroectoderm (NC) (N=292, p<5x10^-4^), and the mesoderm (MC) (N=739, p<5x10^-4^). This supports that *Raf^926^* mutant embryos are transcriptionally similar to the dorsalized embryos that are generated by perturbing the Toll pathway.

Overall, the transcriptional analysis showed that *Raf^926^* mutants recapitulated the effect of strong loss of function mutants in the Toll pathway with most key mesoderm and neuroectoderm determining genes being significantly downregulated and ectoderm determinants being significantly de-repressed. This pattern of expression changes supports that these are dorsalized embryos, which lack both the mesoderm and neuroectoderm germ layers.

### Effect of *Raf^926^* mutants on Dorsal nuclear localization

Based on the lack of a ventral furrow and the loss of Dorsal induced genes, we turned to D/V patterning pathways to determine the mechanism through which Raf is required for ventral furrow formation. The lack of ventral furrow in *Raf^926^* was very similar to *dorsal* and *Toll* mutants, so we evaluated the activity of the Toll pathway in mutant embryos. The Dorsal protein localizes to the nucleus in a dynamic manner in response to Toll signaling in the ventral cells of pre-gastrulation embryos (Fig. 4A, S. Fig. 3A). *Raf^926^* embryos showed a complete exclusion of Dorsal protein from the nucleus in both projection view (S. Fig. 5D-F) and in cross-section view (S. Fig. 5J-L), compared with the expected gradient of nuclear localization seen in wild-type embryos in both projection view (S. Fig. 5A-C) and in cross-section view (S. Fig. 5G-I). This is most readily observed when orienting the embryo with the ventral side centered in the microscope showing a stripe of nuclear Dorsal that is absent in *Raf^926^* embryos (Fig. 4A-F). To observe this process in living embryos, we used Dorsal-GFP to image the early stages of Toll pathway activation. In normal development, a gradient of nuclear Dorsal-GFP can be observed before the ventral furrow forms, and the cells at the center of this gradient invaginate to begin ventral furrow formation (Combined Video 3, Videos 8 and 9, Still Images in S. Fig. 6A - E, S. Fig. 7A – E) [84]. We introduced the Dorsal-GFP construct into *Raf^926^* embryos, which showed no discernible nuclear localization on the ventral side (Combined Video 3: Videos 10, 11, 12, Still Images in S. Fig 6A’- E’, S. Fig. 7A’- E’). The lack of nuclear Dorsal protein suggested that the dorsalized phenotype was due to a lack of Toll pathway activity despite many pathway components being upregulated (Fig. 2C).

**Figure 4.**
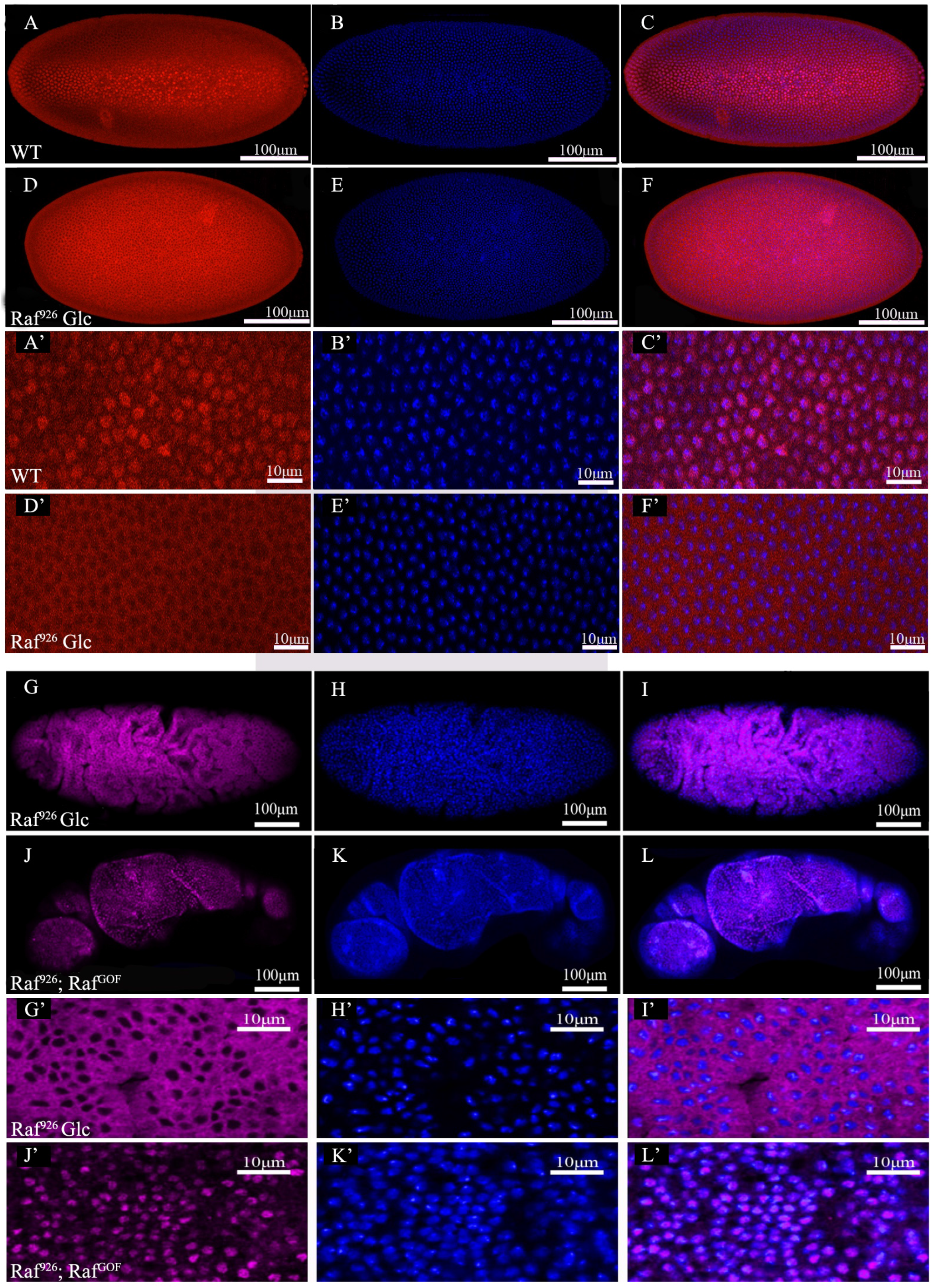
Dorsal nuclear localization in WT, *Raf^926^* embryo and *Raf^926^* embryo expressing Raf^GOF^. Surface view on the ventral side of the **(A, A’)** wild type embryo at cellularization stage showing the stripe of nuclear Dorsal, **(B, B’)** the corresponding nuclear stain and **(C, C’)** merged images of both. **(D, D’)** Complete exclusion of Dorsal from the nucleus in *Raf^926^* embryo at the same stage and view, **(E, E’)** showing the corresponding nuclear stain and **(F, F’)** merged images. **(A’-F’)** were 63x enlarged images of selected regions of **(A-F)**. Surface view of **(G)** *Raf^926^* embryo showing abnormality in development after cellularization, **(G’)** showing the complete exclusion of Dorsal nuclear localization, **(H, H’)** showing the corresponding nuclear stain and **(I, I’)** merged images. **(J)** *Raf^926^* embryo expressing Raf^GOF^ showing partial rescue of the dorsalized phenotype with ventral denticles being restored and **(J’)** showing dorsal nuclear localization. **(K, K’)** shows the corresponding nuclear stain and **(L, L’)** the merged image. **(G’-L’)** were higher magnification images of selected regions of **(G-L)**.

As Raf is not believed to be a Toll pathway component, this finding was unexpected, and suggested genetically that a functional Raf allele was required for ventral Toll pathway activation. This led to a subsequent question: was it also sufficient? We expressed a gain of function Raf transgene in *Raf^926^* embryos to investigate whether Raf alone could induce Dorsal nuclear localization. We observed that expression of Raf^GOF^ (Raf with a constitutively active kinase domain [40]) led to ubiquitous nuclear Dorsal protein in all cells of the gastrulating embryo (Fig. 4J-L), shown in close-up (Fig. 4J’-L’). No nuclear

Dorsal was observed in gastrulating *Raf^926^* embryos not expressing Raf^GOF^ (Fig. 4G-I and in close-up G’- I’). The effect was difficult to observe in pre-gastrulation embryos due to the delay in expression of the GAL4/UAS system, but in later stages the data suggested Raf activity was sufficient to induce Dorsal nuclear localization.

### Raf is Required for Twist expression and Mesoderm development

Ventral furrow cells form the mesoderm after invagination. This process requires an epithelial to mesenchymal transition which is mediated by the *twist* and *snail* genes [85, 86]. Twist is first expressed within the presumptive mesoderm located on the ventral side of the embryo [86–89]. It acts with Dorsal to establish the presumptive mesoderm, and once mesoderm differentiation begins, Twist expression is maintained. Twist and Snail then drive mesoderm formation [89, 90]. In our transcriptional analysis we observed that the *Raf^926^* mutation showed a significant decrease in the expression of both (Fig. 3A).

To fully characterize the effect of Raf on ventral furrow formation, we next looked at the expression of Twist using a live imaging approach. Since the mesoderm is established by ventral cells, Twist in this case acts as a ventral mesoderm marker that gives us insight on how abnormal ventral development in early stages can disrupt late developments of the mesoderm. In a wildtype embryo (Combined Video 4: Video 13, Still Images in S. Fig. 8A - D) the ventral cells developed properly during cellularization and gastrulation, leading to ventral furrow formation and normal mesoderm development, as displayed by twist-GFP’s patterning of the late mesoderm. By contrast, *Raf^926^* embryos lacked twist-GFP signal showing only yolk autofluorescence (Combined Video 4: Videos 14, 15, Still Images in S. Fig. 8A’ - D’).

### Raf signals downstream of Toll

The previous experiments placed the effect of *Raf^926^* upstream of Dorsal nuclear localization but did not establish at what point the Toll pathway was affected. To test this, we expressed a gain of function version of Toll (Toll^10b^) in *Raf^926^* mutant embryos. Toll^10b^, when expressed ubiquitously, activates the signaling pathway in a ligand independent manner resulting in ubiquitous Dorsal nuclear localization and ventralized embryos [3]. As an intracellular kinase, Raf could function downstream of the Toll receptor. We observed that expression of Toll^10b^ had no effect on Dorsal protein localization or on the dorsalized phenotype of *Raf^926^* maternal and zygotic mutants (Fig. 5A, A’, A’’, B, B’, B’’). This was not due to a lack of activity, as paternally rescued *Raf^926^* embryos did show some ventral tissue restored (S. Fig. 9), but Toll^10b^ had no effect on maternally and paternally mutant *Raf^926^* embryos, genetically placing *Raf^926^* downstream of Toll (S. Fig. 10B).

**Figure 5.**
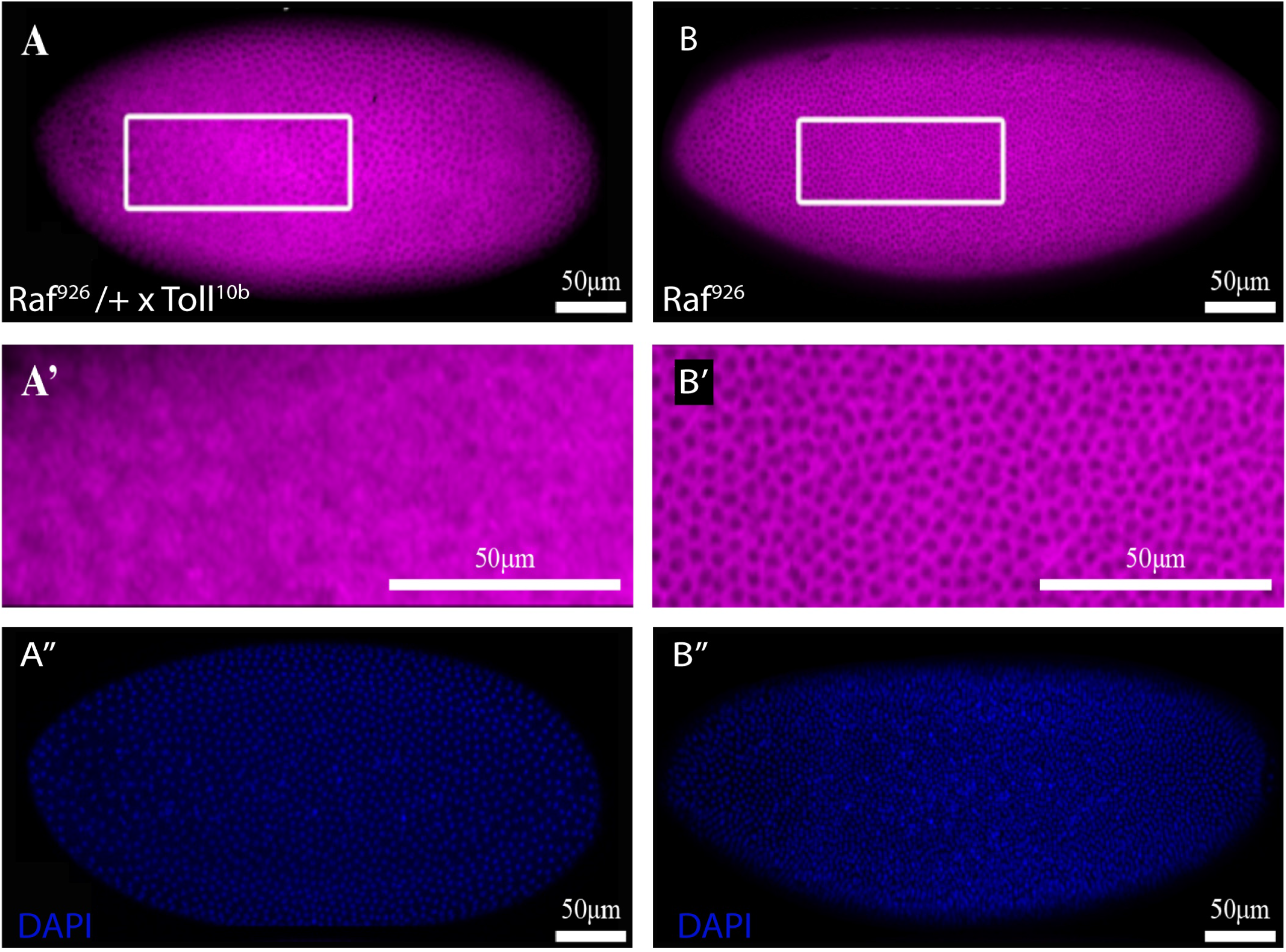
Dorsal nuclear localization in *Raf^926^* embryo expressing the gain-of-function *Toll^10B^*. Surface view of **(A)** paternally rescued *Raf^926^* embryo expressing *Toll^10B^* showing small degree of nuclear Dorsal localization. **(A’)** enlarged image of the boxed area in **(A)**, showing partial dorsal nuclear localization towards the anterior end of the embryo. **(B)** *Raf^926^* embryo with a higher magnification view **(B’)** showing complete dorsal nuclear exclusion. **(A’’ and B’’)** Nuclear stain of corresponding embryos.

We extended the epistasis analysis by investigating whether an activated Ras, Ras^V12^, which functions upstream of Raf in the EGF pathway, could rescue the *Raf^926^* phenotype. As with the activated Toll crosses, Ras^V12^ was unable to rescue the dorsalized phenotype either early or late in development (S. Fig. 10D as compared to S. Fig. 10A). Looking further downstream in the Toll pathway, we expressed RNAi against *cactus* to determine if *Raf^926^* acts upstream of *cactus* in the Toll pathway. We found that embryos homozygous maternally and paternally for *Raf^926^* and expressing RNAi knockdown of *cactus* showed nuclear Dorsal, especially late in development (S. Fig. 10F), producing a similar result as the *Raf^926^* embryos crossed with activated Raf (S. Fig. 10E). These results demonstrated that *Raf^926^* was upstream of Cactus and downstream of Toll and Ras. Nuclear Dorsal could also be observed in embryos overexpressing Dorsal (S. Fig. 10-5C).

To investigate the molecular mechanism through which Raf acts on the Toll pathway more closely, we then turned to a cell culture model, where we first verified our *in vivo* findings in a tightly controlled environment. We designed two signaling sensors, the traditional MAPK cascade target ERK fused to RFP as the control for EGF signaling, and Dorsal fused to RFP as the indicator for Toll signaling. Expression of an activated Ras^V12^ was sufficient to induce both Dorsal and ERK nuclear localization. Expression of Raf^926^ did not induce nuclear localization of either Dorsal or ERK (Fig. 6A-D).

**Figure 6.**
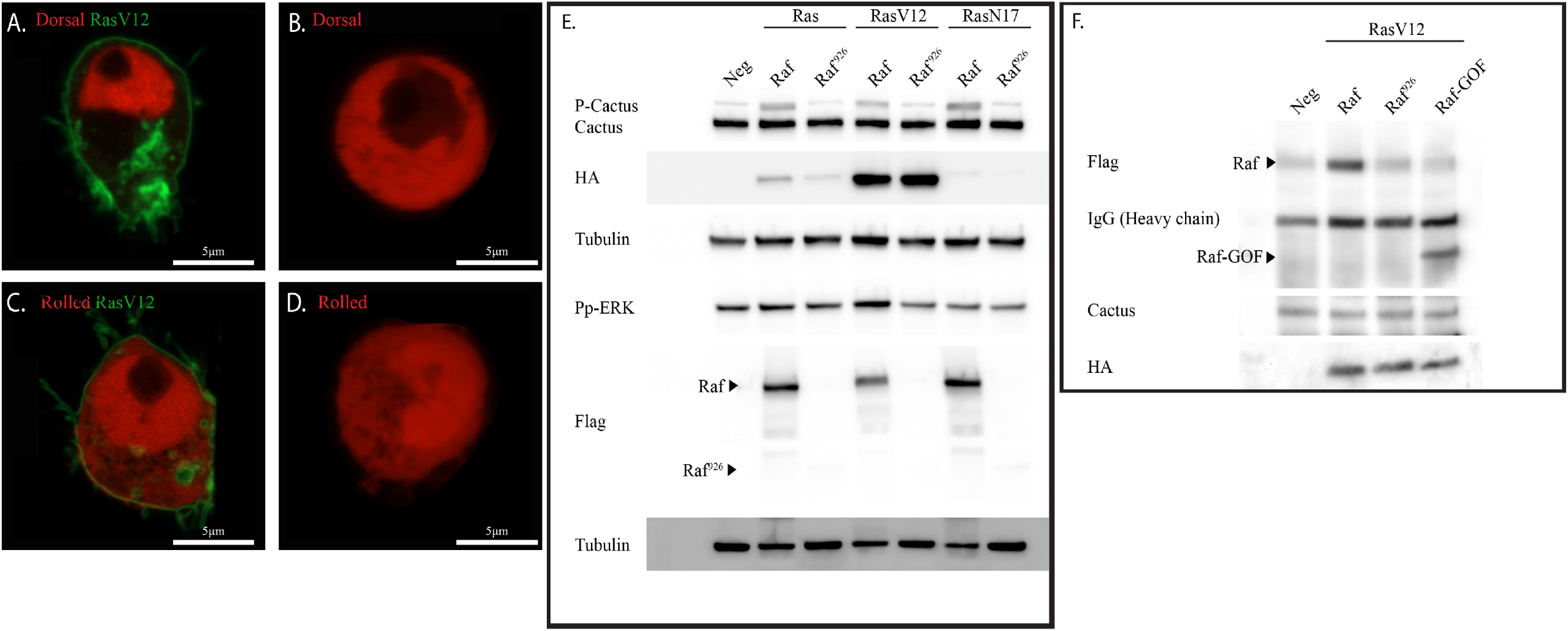
Dorsal and MAPK localization in vitro. **(A)** Nuclear localization of Dorsal (red) in S2R+ cell expressing the gain of function Ras^V12^ (green). **(B)** Nuclear exclusion of Dorsal in S2R+ cell in the absence of RasV12 expression. Ubiquitous expression of Erk (red) in both cytoplasm and nucleus of S2R+ cell both **(C)** in the presence and **(D)** absence of Ras^V12^ (green). The black spot in the nucleus in **(A-C)** is the nucleolus. **(E)** Protein levels of phosho-Cactus (P-cactus), Cactus and Pp-ERK in S2R+ cells transfected with flag-tagged Raf or Raf^926^ and HA-tagged Ras, gain of function Ras^V12^ or dominant negative variant Ras^N17^. Higher phosphorylated cactus (P-cactus) level observed in cells transfected with Raf compared to Raf^926^ and the negative control. Untransfected S2R+ cells served as negative control, and tubulin as loading control. The deletion-frameshift mutation in Raf^926^ resulted in nonsense decay of flag- Raf^926^, rendering it undetectable. Triangle indicates the expected molecular weight of flag- Raf^926^ in the immunoblot. **(F)** Co-immunoprecipitation of flag-Raf and HA-Ras pulled down with Cactus antibody. Immunoblot showing an interaction between Raf and Cactus in S2R+ cells transfected with either flag-tagged Raf, Raf^926^ or Raf^GOF^ and HA-tagged RasV12.

Previously, studies into the oncogenic properties of Raf showed that, in mammalian cell culture, Raf regulates NFκB signaling directly by binding and phosphorylating IκB [91]. We therefore sought to determine whether *Drosophila* Raf was binding and phosphorylating the IκB homologue Cactus. In cells transfected with active Raf or Ras, we observed an increase in Cactus phosphorylation as visualized by a higher migrating band compared to negative controls or cells transfected with the Raf-926 construct (Fig. 6E) [92]. We also observed an interaction between Raf and Cactus in co-immuno precipitation experiments (Fig. 6F). Taken together with our results showing that Cactus RNAi can rescue the *Raf^926^* phenotype, these experiments suggest that much like human Raf, *Drosophila* Raf may play a role in Toll signaling during the establishment of dorsal-ventral polarity in embryos.

### Early embryonic EGF activation through Optogenetics

As seen in the Raf^GOF^ experiments, activating EGF signaling in very early stages of embryogenesis was not possible using the standard Gal4/UAS system. We took advantage of a recently developed model of EGF signaling which uses an optogenetically activatable allele of SOS [93]. Opto-SOS has previously been used to study patterning, transcription and morphogenesis [94, 95]. A major finding of this work was that hyper- activation of the EGF pathway led to fate switching depending on cumulative ERK activity. Compressive forces moved yolk within the gastrulating embryo leading to popping, or ejection of yolk from the interior of the embryo to the exterior [94]. A major advantage of Opto-SOS is that expression of the allele alone does not affect flies if they are kept in the dark, so expression can be driven maternally in the F2 generation leading to very early EGF activation. We used a combination of Gal4 drivers combining a maternal tubulin driver with a constitutive zygotic driver (matαTub-Gal4VP16 and Daughterless-Gal4) to drive early expression of Opto-SOS. We found that *Raf^926^* embryos also showed yolk ejection to the exterior making these embryos very difficult to image at later stages (Combined Video 5, Videos 16-21), and required special preparation for cuticle preparations to remove the unused yolk (Fig. 1). Taking the gain-of-function and loss-of-function findings together suggests that ERK activity is required for a variety of force generating cellular processes during early gastrulation.

The loss of Raf led to a loss of ventral cell fates and to a loss of Twist expression both in transcription (Fig. 3B) and live imaging (Fig. 7A). Activation of Raf through expression of OptoSOS can also inhibit ventral furrow formation through activation of *huckebein* and *WntD* [96], which we observed to be strongly downregulated in *Raf^926^* embryos (Fold Expression Change log2 -6.00 and -8.95 respectively, Supplementary Table 1). In contrast to the loss of function condition (S. Fig. 8), OptoSOS expression activated Twist-GFP expression in all cells of the ectoderm (Fig. 7B). These findings taken together with [96] suggest that Raf hyper activation and loss both affect ventral furrow formation.

**Figure 7.**
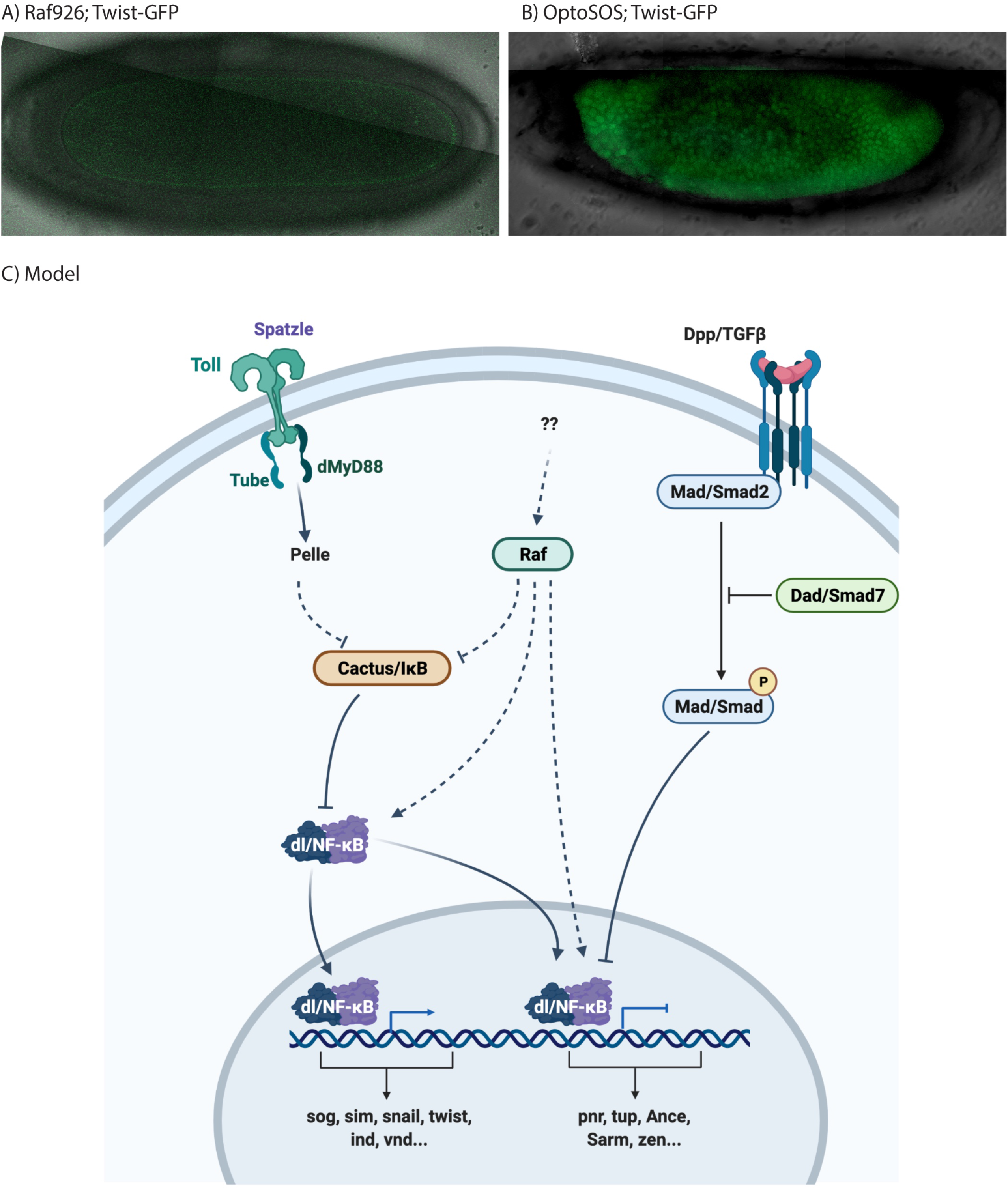
Opto-SOS drives Twist nuclear localization. **(A)** Live, confocal imaging of Twist-GFP expression in a Raf^926^ embryo shows no expression. **(B)** Activation of Raf using Opto-SOS showed uniform Twist-GFP **(C)**. A model for Raf in Dorsal/Ventral signaling.

## Discussion

Previous studies have revealed the multiple roles of Raf in developing the body axes of *Drosophila*. During oogenesis, Raf functions through Pipe in establishing dorsoventral fates of the egg. Targeted Raf activation in follicle cells is sufficient to dorsalize the eggshell whereas reduced Raf ventralizes the eggshell (Brand and Perrimon, 1994). During embryogenesis, Raf defines terminal structures and establishes ventral fates of the neuroectoderm. Early maternal/zygotic mutant screens identified Raf (*pole hole*) mutants as lacking terminal structures and lacking visible ventral cuticle especially in amorphic alleles inducing mutations in conserved region 1 (Fig. 1A) [17, 18, 64]. Although previous studies have comprehensively screened and characterized many Raf mutants, no connection between Raf and the dorsoventral-determining Toll pathway has been shown during embryonic development. We applied a modern screening approach, where a series of mutations were made and characterized through whole genome sequencing [63], and found a strong mutation in Raf that led to a complete loss of ventral structures. We characterized the *Raf^926^* embryo and established its dorsalized phenotype. We discovered that this dorsalized phenotype was not due to defective oogenesis and could not be explained completely by defective *torso* signaling.

Most importantly, this Raf function appears to be independent of the MAPK cascade, as there is no evidence of active MAPK in ventral cells. Studies looking at phosphorylated MAPK show activation at the anterior and posterior poles in early stages, lateral activity in early gastrulation and activation in the mesoderm [80]. Interaction of MAPK and Toll pathway has been observed at early stages through Capicua and WntD [97, 98]. This suggests that Raf’s function in the most ventral cells is not through MAPK, but could perhaps function through direct phosphorylation of IκB/Cactus [99]. Alternatively, Raf could act similarly to MAP3K14, also known as NIK (NF-κB inducing Kinase), which cooperates with IKKα in mammals to phosphorylate IκB, thereby activating NF-κB signal [49, 50, 100]. As Raf is a MAP3K, it could play a similar role in *Drosophila* Toll signalling, but additional investigations into the precise molecular mechanism connecting Raf and the Toll pathway will be required. Alternatively, Raf could function in conjunction with Sterile 20 like kinase in a MAPK-independent, parallel pathway [60, 61].

In short germband insect species, EGF signaling helps to establish dorsoventral polarity [101], and our results suggest a role for the EGF signaling component Raf in the long germband insect *Drosophila*. EGF signaling is involved in patterning of the oocyte where signaling from the egg to the follicle cells establishes egg polarity [30, 102]. In the embryo, EGF’s dorsal-ventral patterning activity is thought to be limited to the neuroectoderm. Both the Toll and EGF pathways are highly conserved in oncogenesis and immune function, and this study presents the first *in vivo* evidence that EGF and Toll signaling are connected in a normal physiological/developmental system [103–106]. Further characterization of signalling events bridging EGF and Dorsal/NF-κB signalling could yield valuable insight into the regulation of the therapeutically important NF-κB family of proteins and broaden our understanding of how neoplasms which have attained EGF-family mutations interface with the body’s immune surveillance mechanisms.

## Methods

### Crosses and expression of UAS construct

Maternally mutant eggs were generated by the dominant female sterile technique where balanced mutants are crossed to the dominant female sterile mutation *Ovo^D2^* and recombination is induced using the FLP/FRT method in ovaries [66, 107]. The *Ovo^D2^*, FRT19A double mutant line was generated by recombining X chromosomes with *Ovo^D2^* and FRT19A in a female rescued for sterility by a second chromosome duplication carrying a wildtype *Ovo* gene (Dp(1;2)w+64b). Oregon R was used as the wild-type strain. Please see Flybase for further details on mutants used (flybase.bio.indiana.edu). Mutants used: *Raf^926^*. For mis-expression experiments, we used a GAL4 driver combining the early expression of the matαTub-Gal4VP16 together with daughterless-Gal4 on the third chromosome (matda- gal4). All X-chromosome mutants use FRT19A. The following crosses were conducted:

1. *w*, *Raf^926^*, FRT19A/ FM6; matda-gal4/+ female x *w*, *ovo^D2^*, FRT19A/male
2. *w*, *Raf^926^*, FRT19A/ *Ovo^D2^*, FRT19A female x FM6/male
3. *w*, *Raf^926^*, FRT19A/ *Ovo^D2^*, FRT19A; matda-gal4/+ female x UAS-Raf^926^-mCherry, attP2
4. *w*, *Raf^926^*, FRT19A/ *Ovo^D2^*, FRT19A; matda-gal4/+ female x UAS-Raf^GOF^ [40]
5. *w*, *Raf^926^*, FRT19A/ *Ovo^D2^*, FRT19A; matda-gal4/+ female x UAS-Ras^V12^ [108]
6. *w*, *Raf^926^*, FRT19A/ *Ovo^D2^*, FRT19A; matda-gal4/+ female x UAS-Toll^10b^ [109]
7. *w*, *Raf^926^*, FRT19A/ *Ovo^D2^*, FRT19A; matda-gal4/+ female x UAS-Cactus TRIP RNAi [110]
8. *w*, *Raf^926^*, FRT19A/ *Ovo^D2^*, FRT19A x Dorsal-GFP [84]
9. *w*, *Raf^926^*, FRT19A/ *Ovo^D2^*, FRT19A x Twist-GFP [111]
10. *w*, *Raf^926^*, FRT19A/ *Ovo^D2^*, FRT19A x QUAS-GFP, Tubulin-QF2 [112]
11. *w*, *Raf^926^*, FRT19A/ *Ovo^D2^*, FRT19A x UAS-Dorsal-Cry2-mCherry [113]
12. *w*, *Raf^926^*, FRT19A/ *Ovo^D2^*, FRT19A x Zcl-GFP [114]
13. *w*, *Raf^A^*, FRT19A/ *Ovo^D2^*, FRT19A [63]
14. *w*, *Raf^B^*, FRT19A/ *Ovo^D2^*, FRT19A [63]
15. nanos-gal4; UAS-Cas9 x Raf TRiP- KO guide RNA [69]
16. matda-gal4 x UAS-OptoSOS [93]
17. matda-gal4 x UAS-OptoSOS; Twist-GFP [111]
18. matda-gal4/+ female x UAS-Raf^926^-mCherry, attP2 (This study)

X chromosomes were marked with *w^1118^* mutation and the CyO and FM6 balancers were marked GFP to simplify analysis. As mothers were heterozygous for the Gal4 source, maximal rescue is reflected by a drop of phenotype to 50% (only half of the embryos will express Gal4). For all crosses, more than 100 embryos were analyzed in multiple, separate experiments (n>95). Additional stocks were obtained from the Bloomington stock center including the Q system [115].

### Mutant, transgene and driver lines

The mutation was genetically mapped on the chromosome X by complementation with deletion strains, and precisely localized to 3A1-A2 through suppression by the molecularly defined P[acman] duplication DC107 which covers Raf’s protein coding exons or the RA splice variant, but not the non-coding regions in the RE splice variant [116]. The molecular nature of the *Raf^926^* allele with a deletion of 17 nucleotides (GAGTACGACTATGTGAT) from the 479^th^ nucleotide of the protein-coding region, was found by Sanger sequencing. The genomic DNA was amplified by PCR, sequenced, and the products were cloned into pENTR vectors (Invitrogen) and recombined using Gateway technology (Invitrogen) into pUASg.AttB.mCherry vectors for fly injection [117, 118]. The DNA was injected into strains P[CaryP]attP2 68A4 and P[CaryP]attP40 25C6 by BestGene Inc California [119]. For constitutive expression in S2R + cells, the pDONR vectors were recombined into the Gateway destination vector, pAW (Drosophila Gateway® Vector collection, Carnegie Institution), with NH2 terminus flag-tag for Raf, Raf^926^ and Raf^GOF^, HA-tag for Ras, Ras^V12^ and Ras^N17^ and COOH-terminus RFP and GPF-tags for Erk, dorsal and Ras^V12^ constructs used in live imaging. The Toll and Dorsal optogenetic constructs were obtained by gene synthesis and combined with cryptochrome 2 and mCherry cDNA as described in [120].

Additional genetic testing was done with two further novel Raf point mutants, Raf^A^ (H546Y) and Raf^B^ (L163R) obtained from the Bloomington Drosophila Stock Center. CRISPR/Cas9 studies were carried out using Raf TKO.GS00615 gRNA line in combination with a *nanos* driven Cas9 [69].

### Immunofluorescence

Embryos were fixed with Heat-Methanol treatment [121] or with heptane/4% formaldehyde in phosphate buffer (0.1M NaPO4 pH 7.4) [122]. The antibodies used were: 1:10 – 1:100 anti-dorsal (mouse mAb, 7A4, Developmental Studies Hybridoma Bank (DSHB) developed under the auspices of the NICHD and maintained by The University of Iowa, Department of Biological Sciences, Iowa City, IA 52242), 1:500 Hoechst 33342 for nuclear staining. Staining, detection and image processing as described in [123].

### Light-sheet microscopy

Embryos at cellularization stage and earlier (Stage 1-5) were selected using halocarbon oil (Sigma). Embryos were carefully dechorionated using bleach, rinsed twice with water and dried, prior to loading into a capillary filled with 1% low-melting agarose (Sigma). Embryo(s) were then carefully oriented under a dissecting microscope using a thin piece of wire and metal probe such that the embryo was upright with the anterior-posterior axis aligned with the axis or perpendicular to the axis of the glass capillary. The agarose was pushed out of the capillary and the sample was suspended freely in the water-filled sample chamber of the Lightsheet Z1 microscope and imaged with a water immersion objective at 20x.

### Scanning electron microscopy

Formaldehyde fixed embryos (with slight adjustment to the cited method: overnight at 4⁰C with rocking; 8 times post-fixation methanol washes) were washed once and re-hydrated with phosphate buffer for 10 minutes with rocking. Embryos were then applied to a microscopy slide. Phosphate buffer on microscopy slide was removed as much as possible. The slide with embryos was then dried prior to imaging. Imaging was performed with Hitachi TM3030Plus table-top scanning electron microscope at 1000x.

### RNA preparation and RNA-sequencing

Mixed stage embryos, 0 to 24 hours after deposition at 25°C, were dechorionated in bleach, washed in water and 100% ethanol prior to RNA extraction using the ISOLATE II RNA Mini Kit’s protocol (Bioline, UK). The extracted RNA was quantified using Nanodrop (Thermo Fisher Scientific). Library preparation was performed using 1µg of total RNA and sequencing was performed using Illumina HiSeq 4000 System (2 × 151 bp read length, 40 million reads per sample) by NovogeneAIT Genomics (Singapore).

### RNA-seq analysis

#### Data processing and QC

RNA-seq was aligned against BDGP6.22 (Ensembl version 97) using STAR v2.7.1a [124] and quantified using RSEM v1.3.1 [125]. Reads annotated as rRNA, snoRNA, or snRNA were removed. Genes which have less than 10 reads mapping on average over all samples were also removed. Differential expression analysis was performed using DESeq2 [126]. Pairwise comparisons were performed using a Wald test, with independent filtering. To control for false positives due to multiple comparisons in the genome-wide differential expression analysis, the false discovery rate (FDR) was computed using the Benjamini-Hochberg procedure. For the re-analysis of the RNA-seq dataset from Koenecke *et al.*, all of the above is the same except the version of STAR used was v2.9.1a [124]. The gene-level counts were transformed using a variance-stabilizing transformation, converted to z-scores, and clustered using k-means. The total within-cluster sum of squares distances and the elbow criterion was used to determine the optimal number of clusters (k = 3) in this dataset. The significance of the overlap between the RNA-seq data from Koenecke *et al.* and our dataset was computed using random sampling without replacement. After randomly sampling the respective groups of genes, the numbers of overlaps between the two datasets were plotted on a histogram to visualize the probability distribution. The observed number of overlaps was then compared to this distribution and a p-value was calculated.

#### Functional Enrichment analysis

For the analysis of genes upregulated and downregulated in Raf926 mutants (Figure 2c), Gene Ontology (GO) and KEGG pathway enrichments were performed using EnrichR [127–129]. The GO term and KEGG pathway enrichments of the germ layer clusters from the Koenecke *et al*. RNA-seq dataset were also performed with EnrichR. Terms with an FDR < 10% were defined as significantly enriched. The gene sets used in the gene set enrichment analysis (GSEA) were obtained from MSigDB [130–132] via the msigdbr package, and the analysis itself was carried out using clusterProfiler [133].

### Transfection and protein quantification

Drosophila S2R+ cells were maintained in Schneider’s Drosophila Medium with 10% Fetal Bovine Serum and 1% Penicillin and Streptomycin (Invitrogen). Cells were plated at 80% confluence on 60mm dish 24 hours prior to transfection with relevant constructs using the Effectene Transfection Reagent (Qiagen) according to manufacturer’s instructions. Untransfected cells were used as negative control.

36 hours later, cell were lysed with 100µl of RIPA buffer (Cell signaling) with 10% of phosphatase inhibitor and 1% of protease inhibitor (Roche) according to manufacturer’s instructions. Cell lysate were centrifuged at 140,000rpm for 10 minutes at 4⁰C and the supernatant was aliquoted in new tubes at -80⁰C prior to subsequent experiments. The amount of protein in each lysate were quantified using the DC Lowry assay (BioRad) according to manufacturer’s instruction.

### Live cell confocal microscopy

S2R+ cells were plated on 35mm glass-bottom dish at 80% confluence and transfected as mentioned. Cells were imaged 24 hours after transfection using the Zeiss LSM800 confocal microscope with 63x oil immersion lens. Cells transfected with either Dorsal or Erk were used as negative controls for respective experiments. For live embryo imaging, embryos were processed as for Lightsheet microscopy and mounted in 1% low melting point agarose on glass bottomed petri dishes. Imaging was done on a Zeiss LSM 800 (Carl Zeiss, Germany) using the following settings: 1% laser power for 488nm; 5% laser power for 561nm. Images were processed using the ZEN 2014 SP1 software (Carl Zeiss, Germany).

### Co-Immunoprecipitation

Cell lysates were pre-cleared by incubating with 20µl of A/G agarose beads (Santa Cruz) for 30 minutes at 4⁰C with rocking to remove unspecific protein binding to the beads. The mixture was then centrifuged at 2,500 rpm for 10mins at 4⁰C to separate the lysate from the beads. The pre-cleared cell lysate was then incubated with 2µl of anti-cactus primary antibody (mouse mAb, 3H12, DSHB) and 20µl of fresh A/G agarose beads overnight at 4⁰C with rocking. The cell-bead mixture was centrifuged and supernatant was stored if necessary. The beads were then washed three times with 100µl of RIPA (Cell signaling) or salt buffer (50mM Tris, pH 7.4, 400mM NaCl, 5mM EDTA, 1% NP40). Beads were resuspended with 1x loading dye and 1x reducing agent (Invitrogen), and all supernatant was loaded onto the SDS-page gel for Western blotting.

### Western blotting

30µg of protein in 1x sample loading dye with β-mercaptoethanol (BioRad) was boiled at 95⁰C for 5 minutes prior to electrophoresis on a 4-20% gradient SDS-PAGE gel (BioRad). Membrane was blocked with 5% skim milk in TBST 1x on rocker for 1 hour, followed by three times 5 minutes wash with TBST 1x prior primary antibodies overnight incubation. Primary antibodies were used at the following concentration in 5% BSA/TBST 1x: 1:500 anti-cactus (mouse mAb, 3H12, DSHB), 1:1000 for anti-HA (rat mAb, 3F10, Roche), anti-flag (mouse mAb, F9291, Sigma), anti-phospho-p42/44 ERK (rabbit pAb, #9101, Cell Signaling) and anti-tubulin (mouse mAb, T9026, Sigma). Secondary-HRP antibodies: goat-anti-mouse, goat-anti-rabbit (ThermoFisher Scientific) and rabbit-anti-rat R9255 (Sigma) were all used at 1:5000 in 1% skim milk/TBST 1x. Signals were developed with SuperSignal^TM^ West Femto Maximum Sensitivity Substrate and captured with the Syngene C6 gel documentation system and GeneSyn software (Syngene). Images were exported as Tiff and image processing was done via ImageJ (NIH).

## Video Legends

**Combined Video 1. Live imaging of wildtype and *Raf^926^* embryos. (Video 1)** Lightsheet imaging showing that Raf^926^ overexpression (double driver: Arm-Gal4, Daughterless-Gal4>UAS- Raf^926^-mCherry) in an otherwise wildtype embryo does not affect embryonic development. **(Video 2)** Lightsheet imaging showing severe defects in *Raf^926^* embryo expressing Tubulin- QF2>QUAS-GFP and Daughterless-Gal4>Myr-Tomato. **(Video 3)** Development is severely affected as shown by Utrophin-GFP and Histone-RFP lightsheet imaging of *Raf^926^* embryo. **(Video 4)** Normal embryonic development as shown by Utrophin-GFP and Histone-RFP lightsheet imaging of an otherwise wildtype embryo.

**Combined Video 2. Brightfield imaging of embryos from cellularization to early gastrulation. (Video 5)** Normal cellularization to ventral furrow development in a wildtype embryo. **(Video 6)** Lack of ventral furrow formation in a *Raf^926^* embryo. (**Video 7)** Ventral furrow formation in a *bicoid, nanos, torsolike* triple mutant embryo, indicating that the absence of anterior to posterior patterning does not affect ventral furrow formation and is different form the *Raf^926^* phenotype.

**Combined Video 3. Confocal imaging of early embryos expressing Dorsal-GFP. (Video 8)** Nuclear localization of Dorsal-GFP on the ventral side of a wildtype embryo expressing Dorsal-GFP. The cells along the ventral midline invaginate to form the ventral furrow. (**Video 9)** Provides a close-up view. (**Video 10)** No discernible nuclear Dorsal-GFP is present in a *Raf^926^* embryo expressing Dorsal-GFP imaged from cellularization suggesting that the Toll pathway was not properly activated. (**Video 11)** Provides a close-up view. (**Video 12)** Brightfield imaging of the embryo in Video 10 to confirm that embryo was developing.

**Combined Video 4. Confocal imaging of late embryos expressing twist-GFP. (Video 13)** The twist-GFP pattern in the wildtype embryo expressing twist-GFP indicates normal mesoderm development as a result of proper early ventral furrow formation. (**Video 14)** The lack of twist-GFP signal in a *Raf^926^* embryo expressing twist-GFP indicates no proper mesoderm development as a result of the lack of early ventral cell fates and lack of ventral furrow formation. (**Video 15)** Brightfield imaging of the embryo in Movie 14 to confirm that embryo was developing.

**Combined Video 5. Confocal and brightfield imaging of later stage *Raf^926^* embryos. (Video 16)** Brightfield imaging showing yolk movements in *Raf^926^*, **(Video 17)** GFP (Tubulin-QF2>QUAS-GFP) imaging of the same embryo and **(Video 16)** the composite. A second embryo with somewhat different yolk movements **(Video 19-21).**

## Supplementary Figures Legends

**Supplementary Figure 1.**
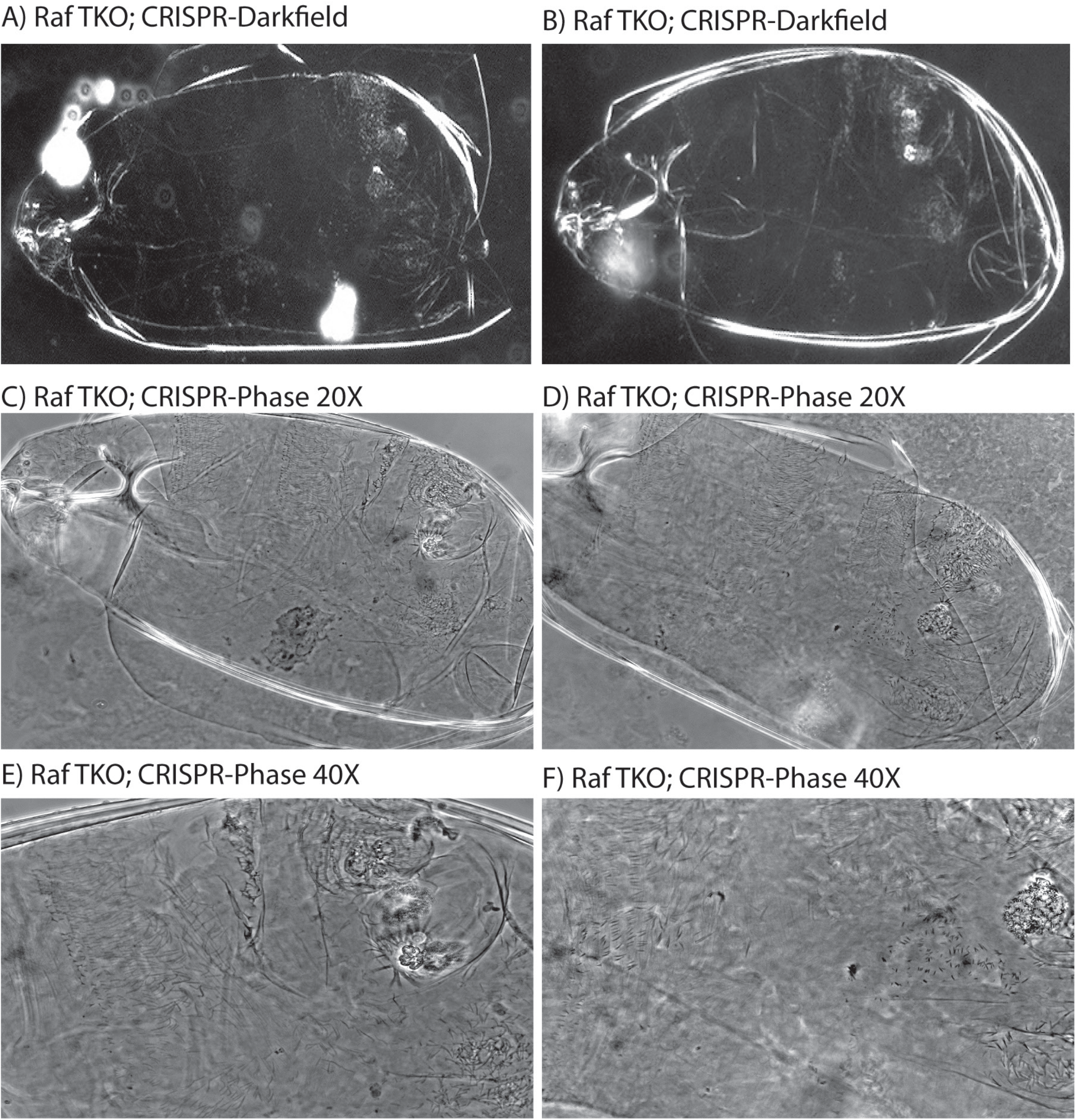
Cuticular preparations of embryos derived from RafTKO.GS00615 gRNA line combined with *nanos-Gal4* driven UAS-Cas9 showing dorsalized embryos. (A) and (B) Embryos imaged using a 10X objective, darkfield. (C) and (D) Embryos imaged using a 20X objective, phase contrast. (E) and (F) Embryos imaged using a 40X objective, phase contrast. All embryos display dorsal hairs and do not produce ventral denticles.

**Supplementary Figure 2.**
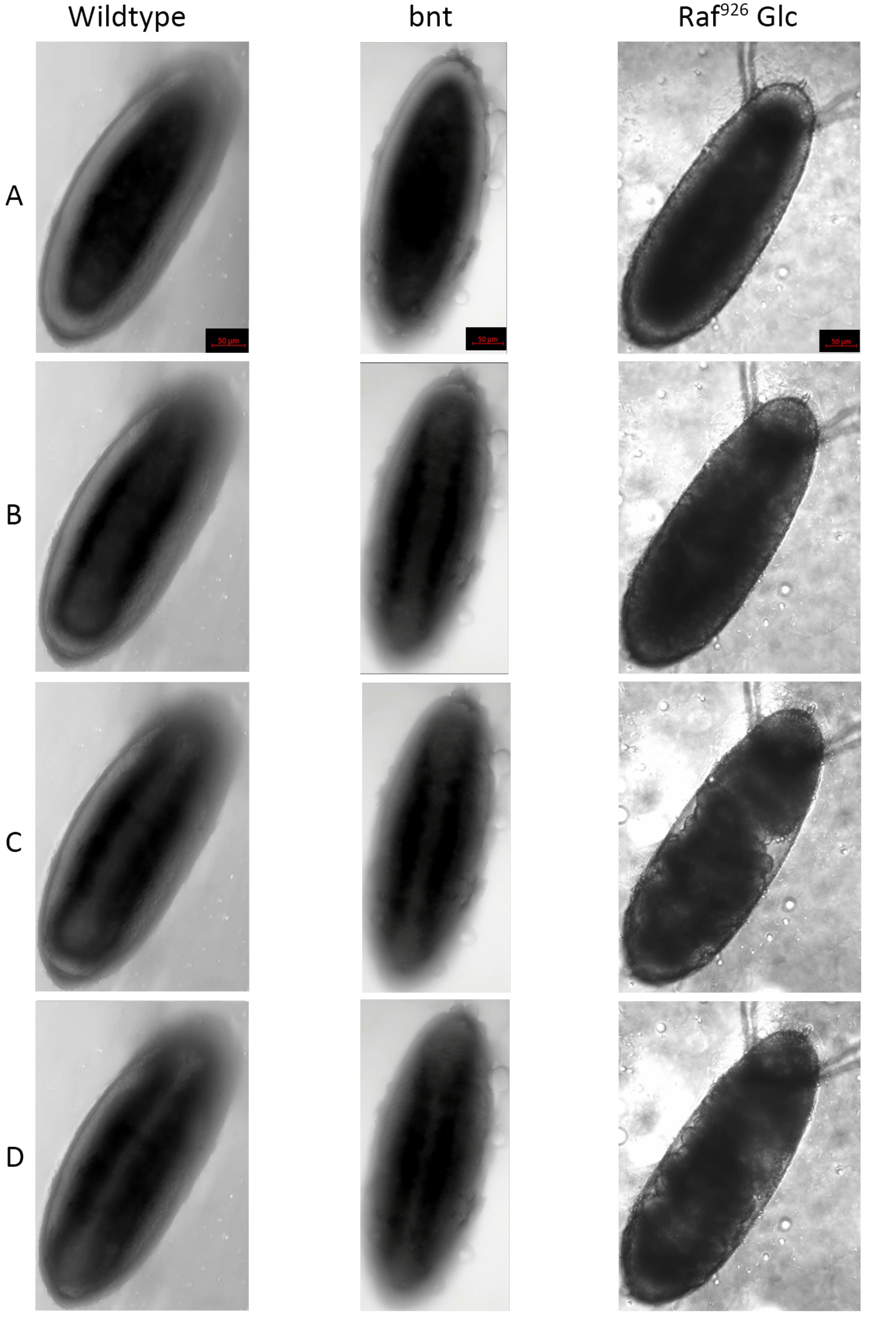
Still images from Videos 5-7 (Combined Video 2). **(A)** shows wildtype, *bnt (bicoid, nanos, torsolike)*, and *Raf^926^* germ-line clone embryos during cellularization. **(B-D)** show progression after cellularization. Wildtype and *bnt* embryos develop ventral furrows whereas *Raf^926^* embryos do not and instead undergo abnormal twisting and turning during gastrulation.

**Supplementary Figure 3:**
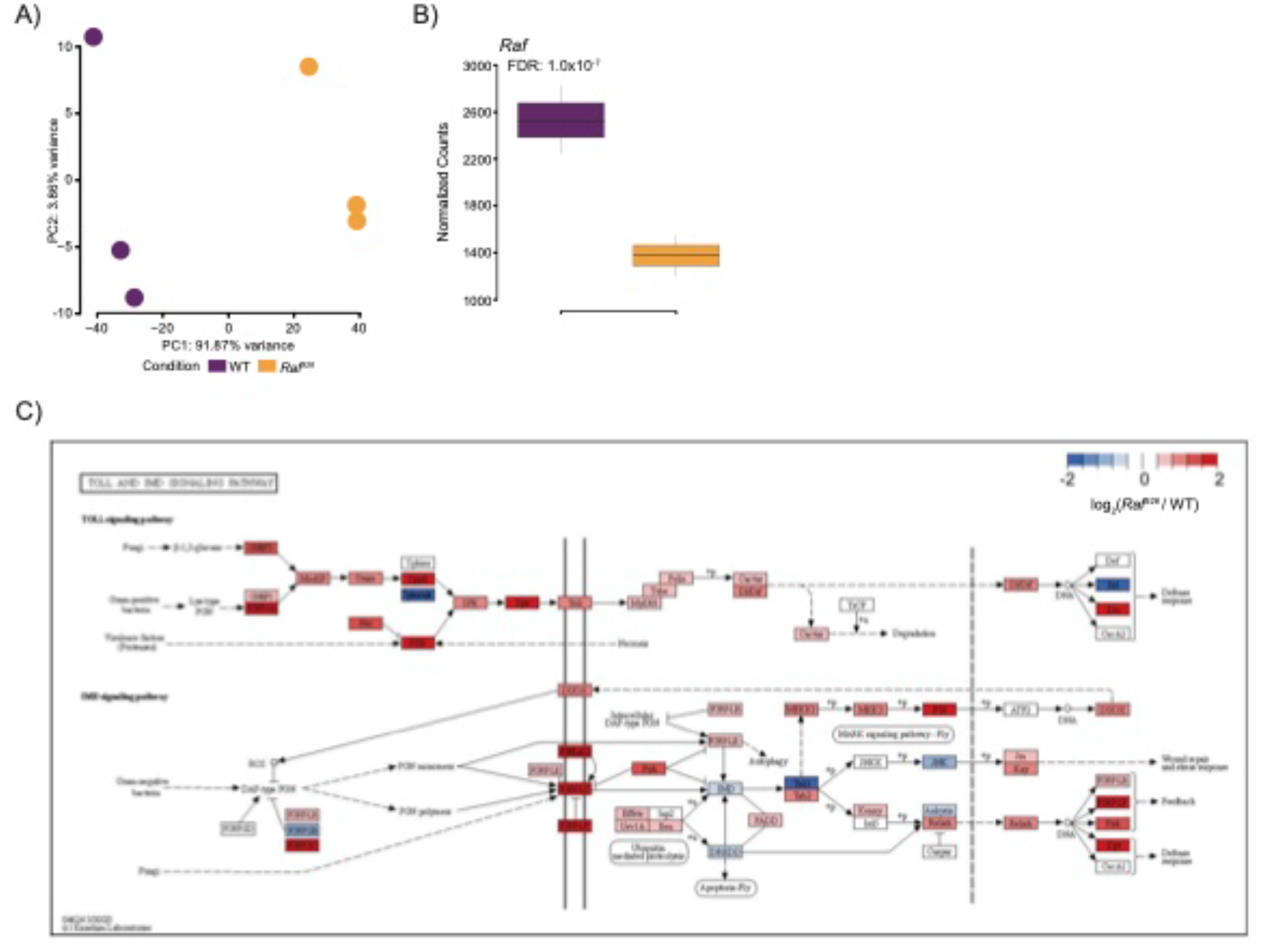
Analysis of RNA-seq from *Raf^926^* mutant embryos. **(A)** Principal component analysis reveals a clear separation between WT and *Raf^926^* embryos, with the first principal component accounting for 92% of the variance. **(B)** Expression of *Raf* is significantly downregulated in *Raf^926^* embryos compared to WT showing less than half the read counts as expected for a maternal effect mutation with heterozygous zygotic expression **(C)** Multiple components of the Toll pathway (KEGG:dme04624) are upregulated in *Raf^926^* mutant compared to WT embryos.

**Supplementary Figure 4:**
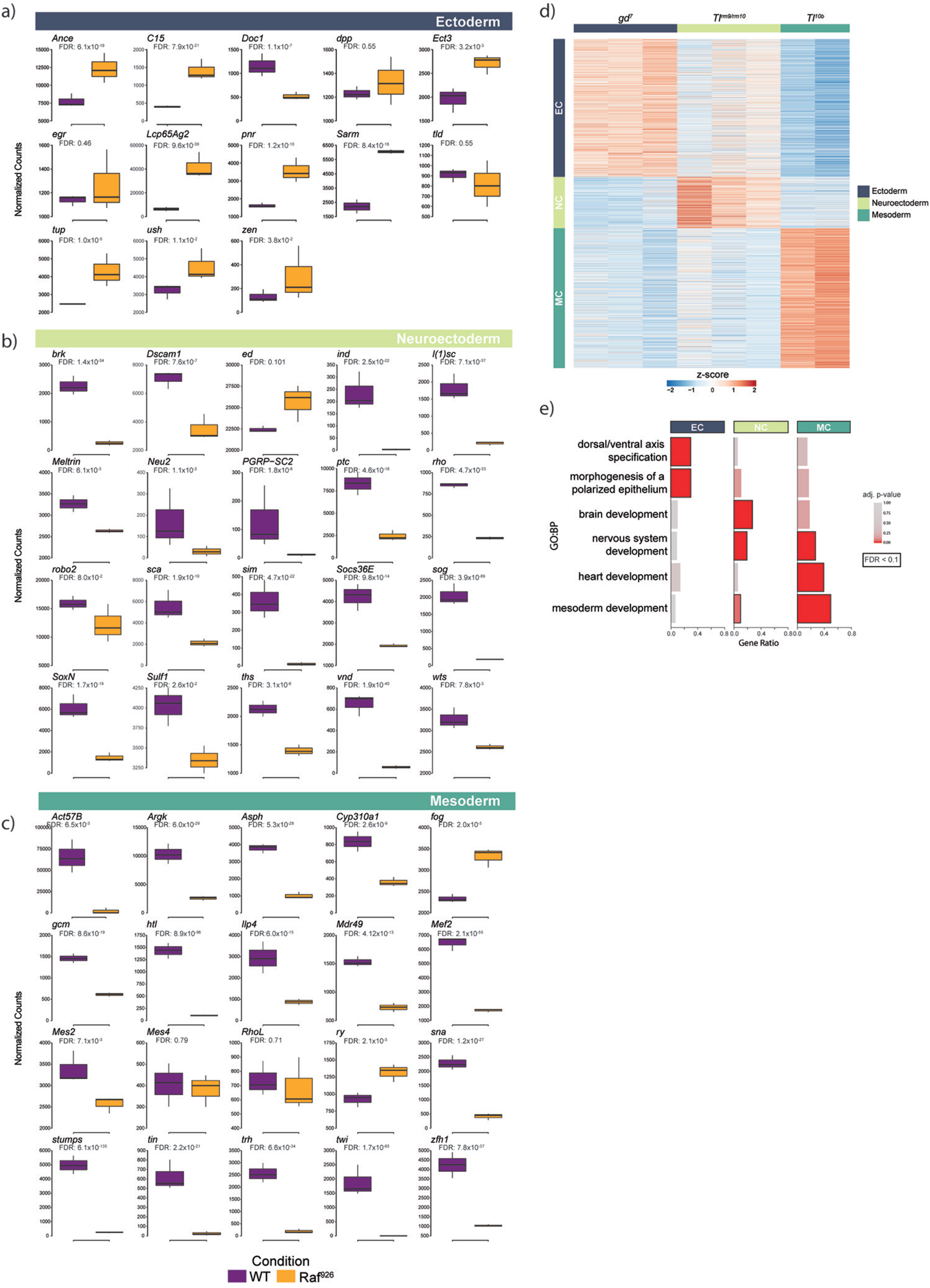
The expression of key germ layer marker genes is affected in *Raf^926^* mutant embryos. Expression changes of all **(A)** ectoderm, **(B)** neuroectoderm, and **(C)** mesoderm marker genes as proposed by [81] that have detectable expression in our model. **(D)** Re-analysis of RNA-seq data profiling from Knoecke *et al.* identifies three clusters which are associated with both distinct mutations in the Toll pathway and distinct germ layers. **(E)** GO Biological process enrichment analysis show that each cluster is enriched for processes associated with the corresponding germ layer (FDR<0.1).

**Supplementary Figure 5.**
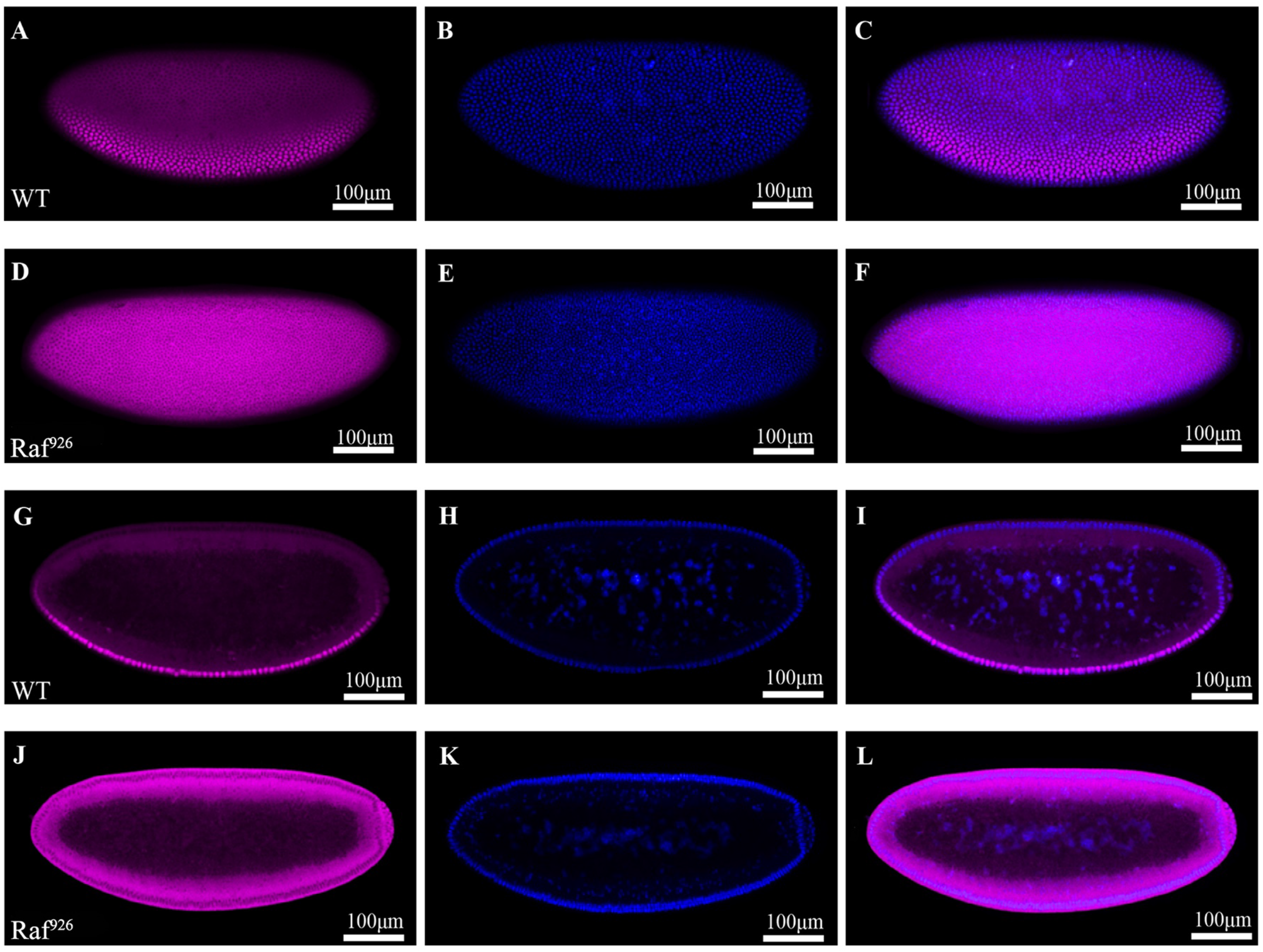
Lack of nuclear Dorsal gradient in the early *Raf^926^* embryo. Surface view of the lateral orientation of the wildtype embryo **(A)** undergoing cellularization with a nuclear Dorsal gradient along the dorsal-ventral axis, **(B)** shows the corresponding nuclear stain and **(C)** a merged image. Surface view of the lateral orientation of the *Raf^926^* embryo **(D)** undergoing cellularization with complete exclusion of Dorsal from the nuclei. **(E)** shows the corresponding nuclear stain and **(F)** a merged image of both. Cross-sectional view of the lateral orientation of the **(G)** wildtype embryo showing the gradient nuclear localization of Dorsal along the dorsal-ventral axis, **(H)** shows the corresponding nuclear stain and **(I)** a merged image of both. Cross-sectional view of *Raf^926^* embryo **(J)** showing the complete exclusion of Dorsal from the nucleus, **(K)** showing the corresponding nuclear stain and **(L)** a merged image of both.

**Supplementary Figure 6.**
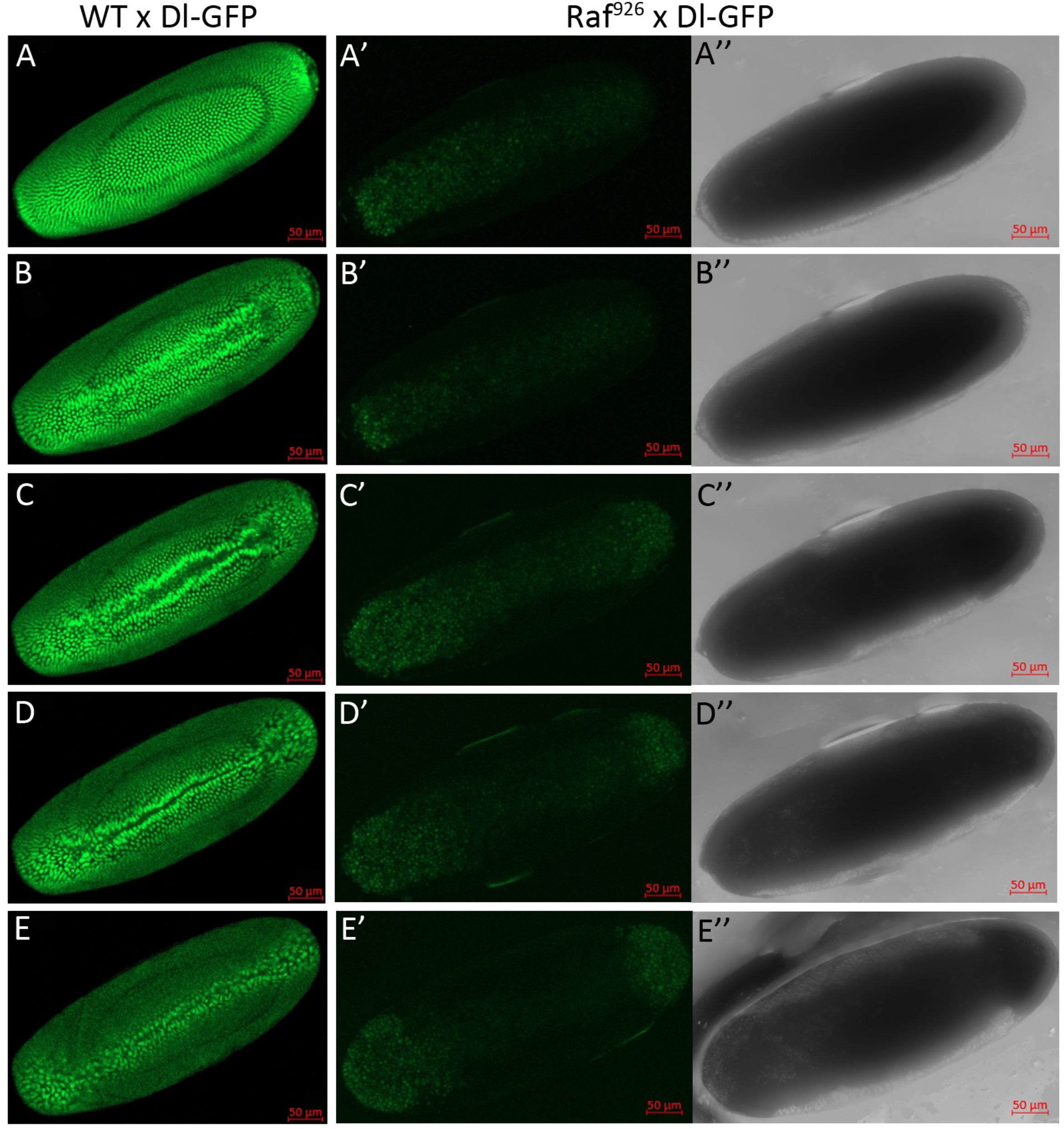
Still images from videos 8, 10, 12 show absence of nuclear Dorsal gradient in live developing *Raf^926^* embryos. (A-E) Surface view of the ventral side of the wildtype embryo expressing Dorsal-GFP from cellularization **(A)** to ventral furrow formation **(B-E)**. Dorsal-GFP is localized in the nucleus. **(A’-E’)** Surface view of the *Raf^926^* embryo expressing Dorsal-GFP from cellularization **(A’)** to early gastrulation **(B’-E’)**, showing no discernible nuclear localization on the ventral side. **(A’’-E’’)** show the corresponding brightfield images of (A’-E’) indicating that the phenotype observed is from a live *Raf^926^* embryo.

**Supplementary Figure 7.**
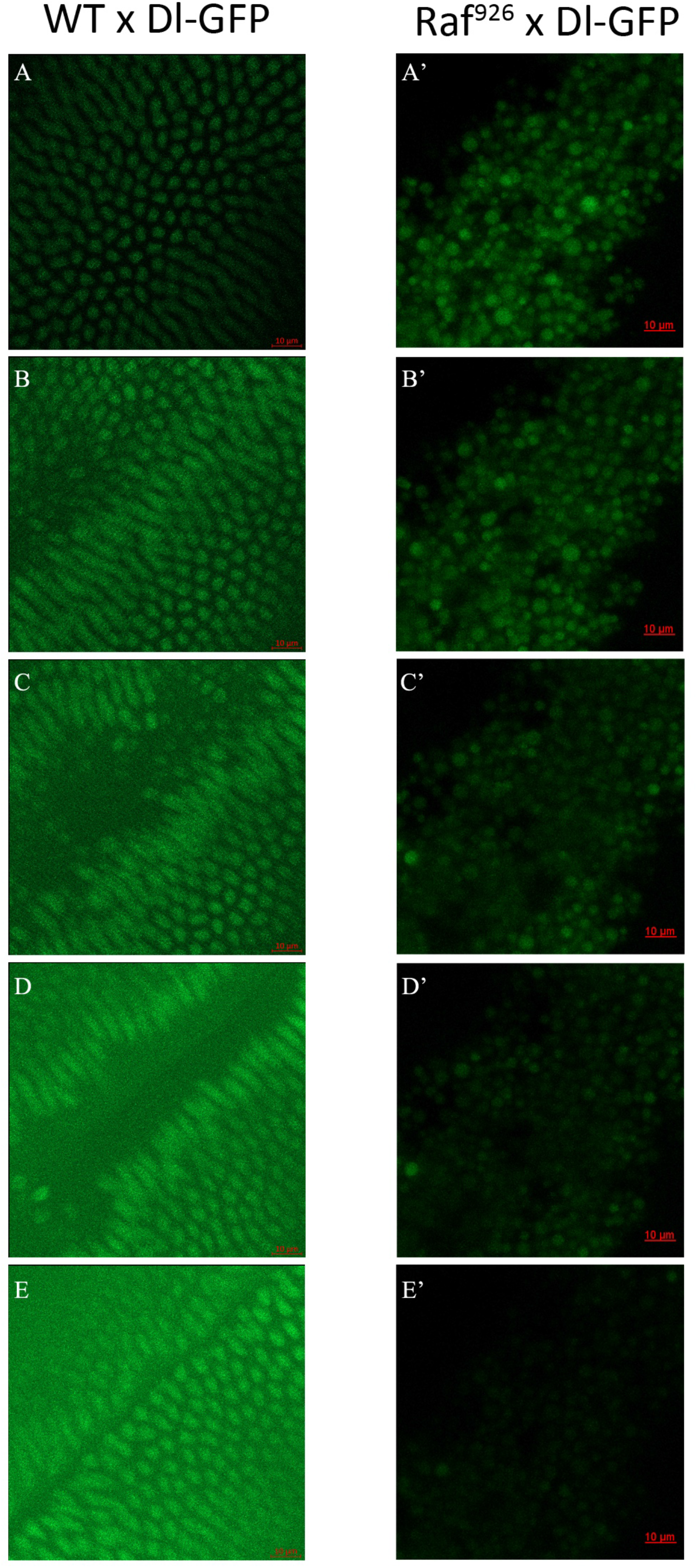
Still images from videos 9 and 11 show no nuclear Dorsal in live developing *Raf^926^* embryos. Close-ups of **(A-E)** surface view of the ventral side of the wildtype embryo expressing Dorsal-GFP from cellularization **(A)** to ventral furrow formation **(B-E)**. Dorsal-GFP is localized in the nucleus. **(A’-E’)** Surface view of the *Raf^926^* embryo expressing Dorsal-GFP from cellularization **(A’)** to early gastrulation **(B’-E’)**, showing no discernible nuclear localization on the ventral side.

**Supplementary Figure 8.**
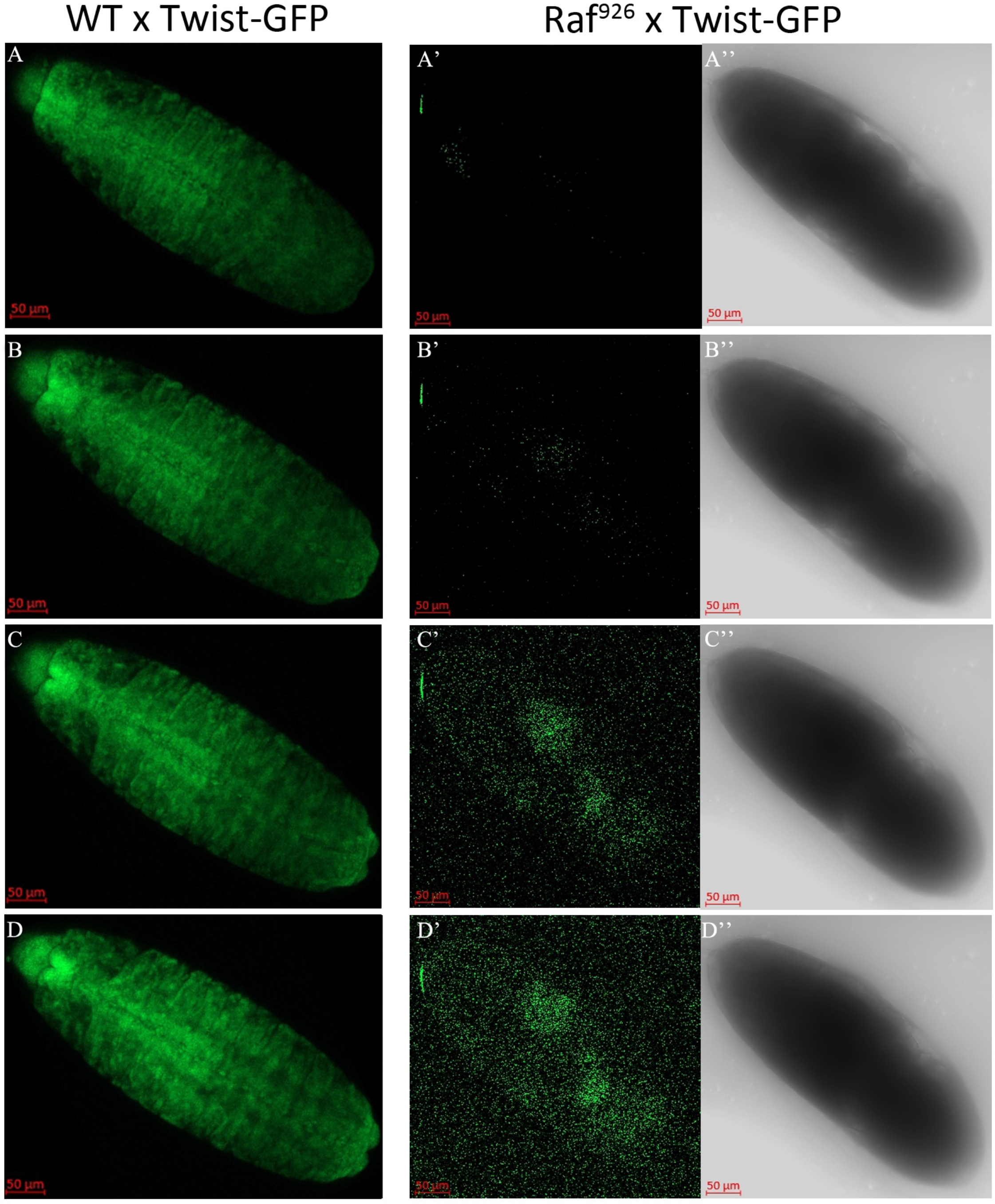
Still images from Videos 13-15 show *Raf^926^* embryos fail to develop mesoderm. (A-D) Late wildtype embryo expressing twist-GFP shows normal mesoderm development. **(A’-D’)** Late *Raf^926^* embryo expressing twist-GFP shows a lack of twist-GFP signal and no proper mesoderm development. **(A’’-D’’)** show the corresponding brightfield images of (A’-D’) indicating that the phenotype observed is from a live *Raf^926^* embryo.

**Supplementary Figure 9.**
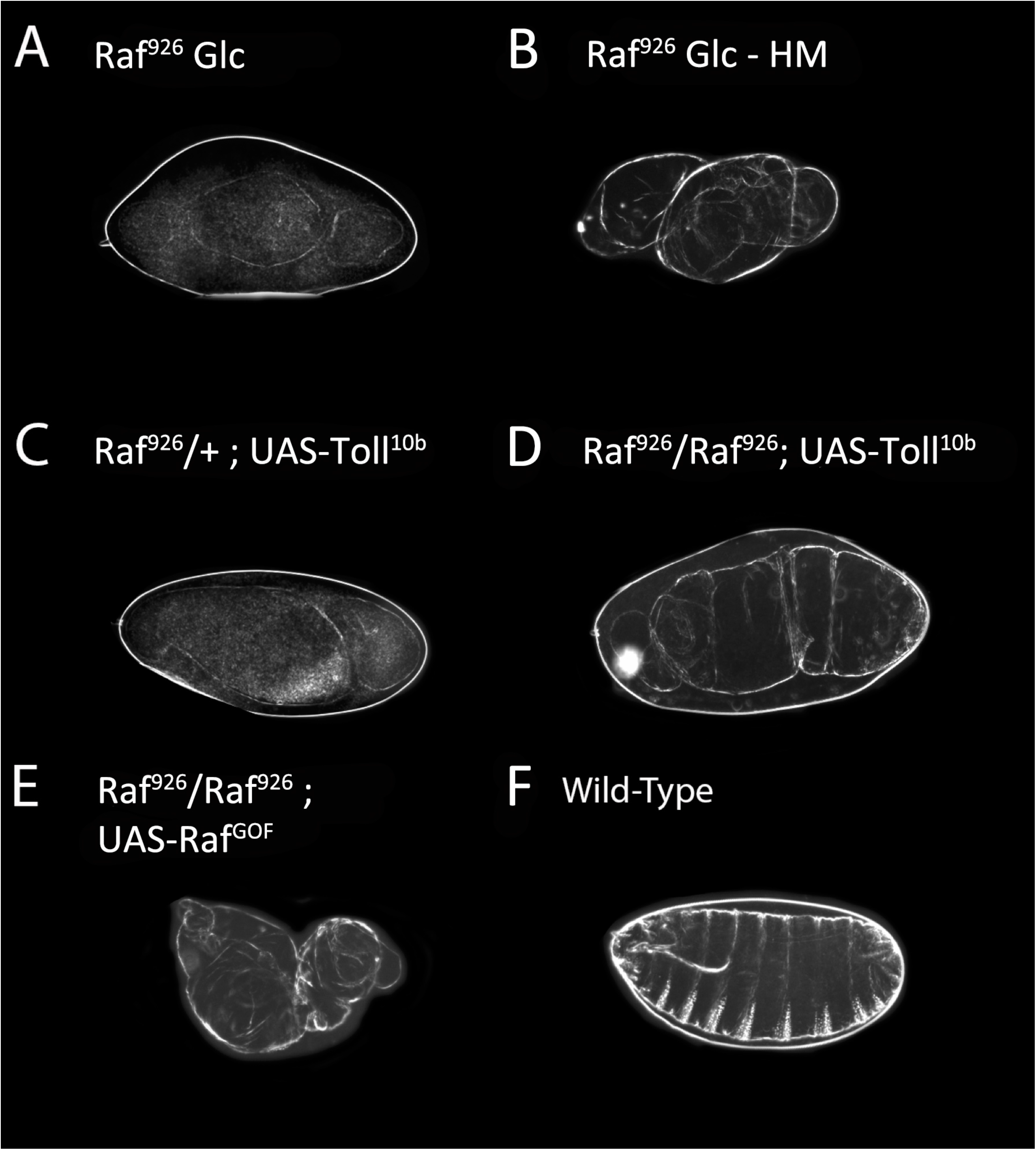
(A) shows a standard cuticular preparation of a *Raf^926^* germ-line clone. **(B)** shows a heat-fixed cuticular preparation of a *Raf^926^* germ-line clone embryo, showing dorsal hairs. **(C)** shows a *Raf^926^* heterozygote expressing UAS-Toll^10b^ while **(D)** shows a Raf^926^ homozygote expressing UAS-Toll^10b^. **(E)** shows a ventralized Raf-Null homozygote expressing UAS-Raf^GOF^. **(F)** shows a wild-type embryo, anterior to the left and ventral to the bottom of the figure.

**Supplementary Figure 10.**
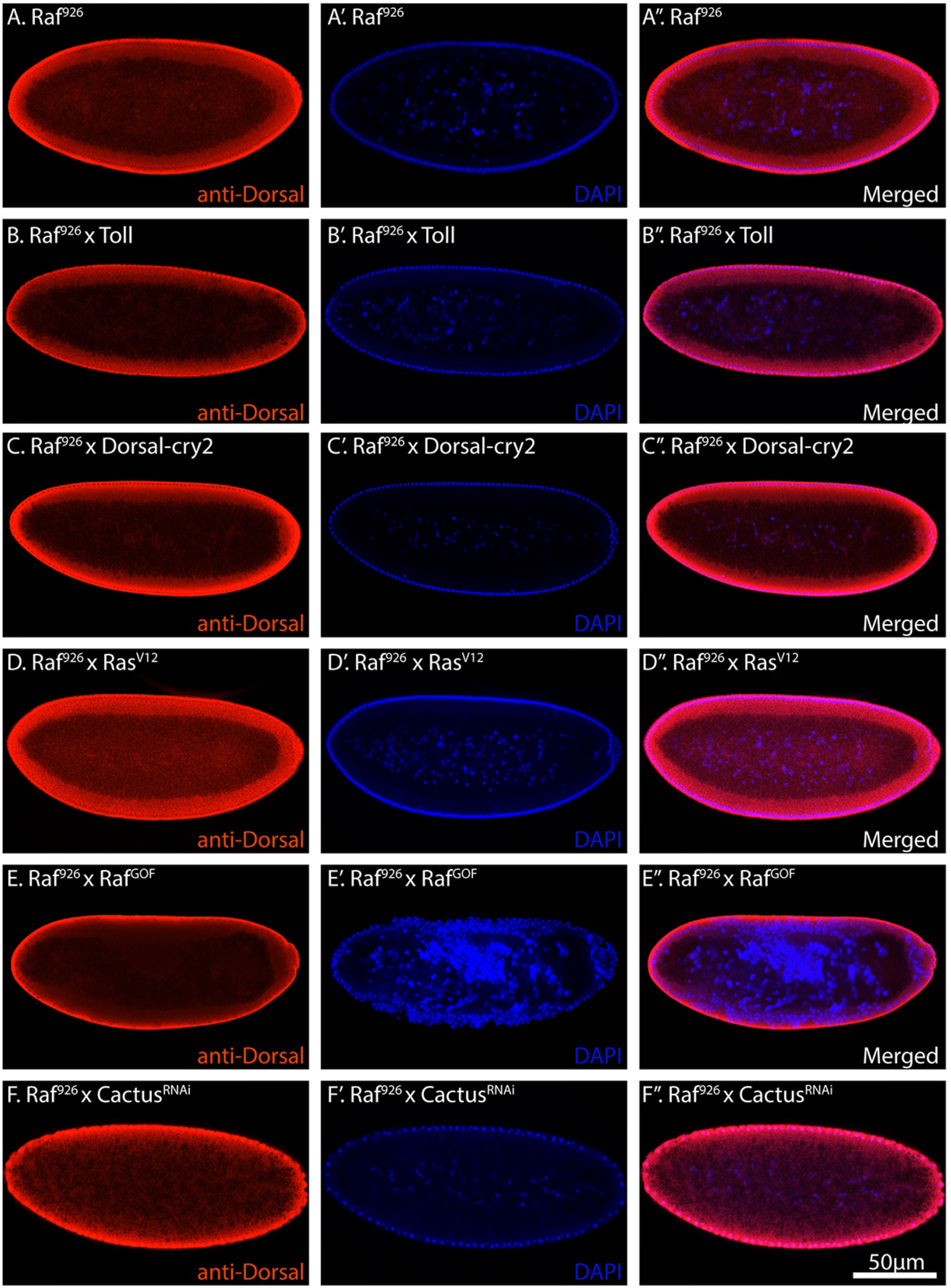
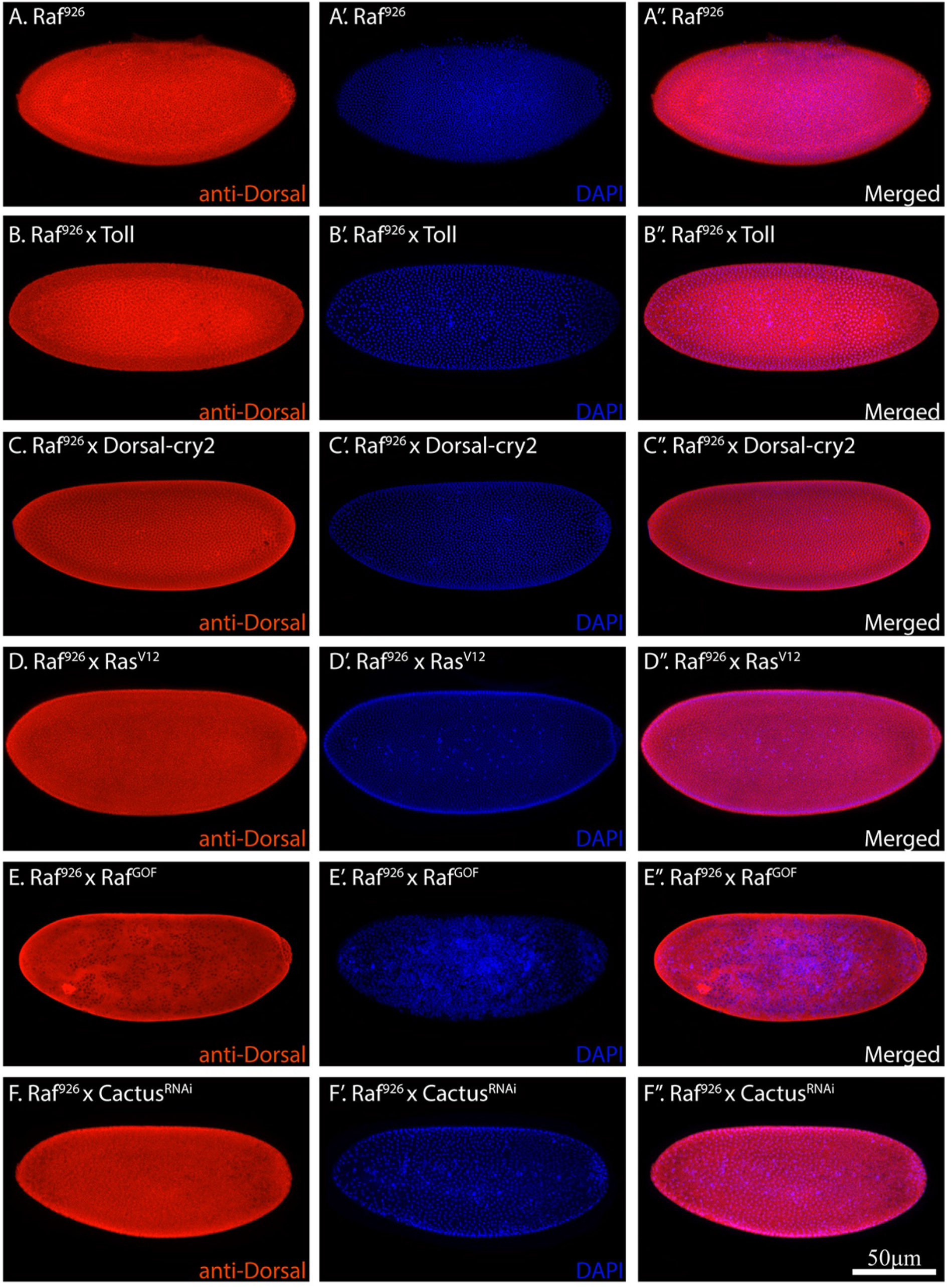
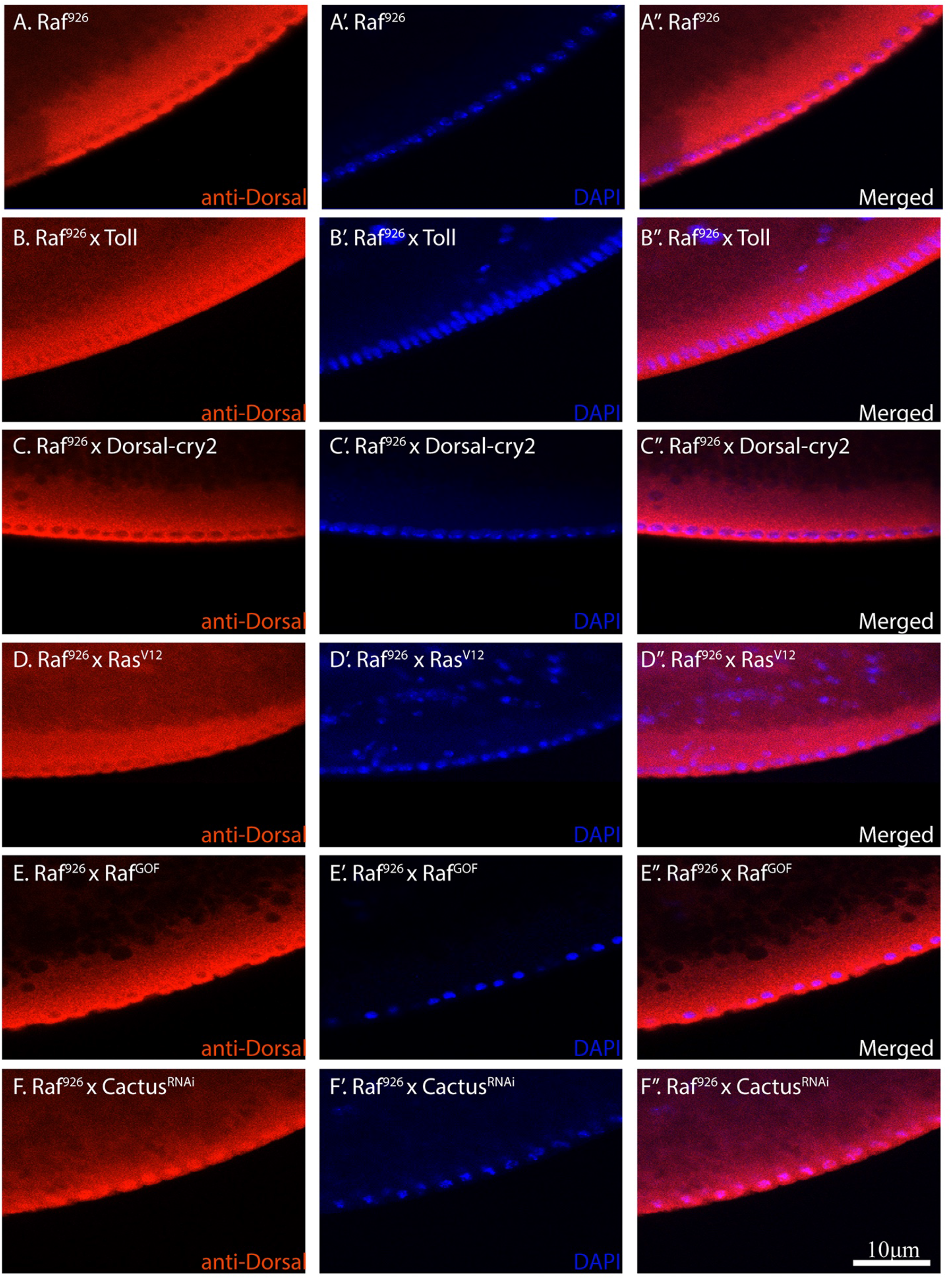
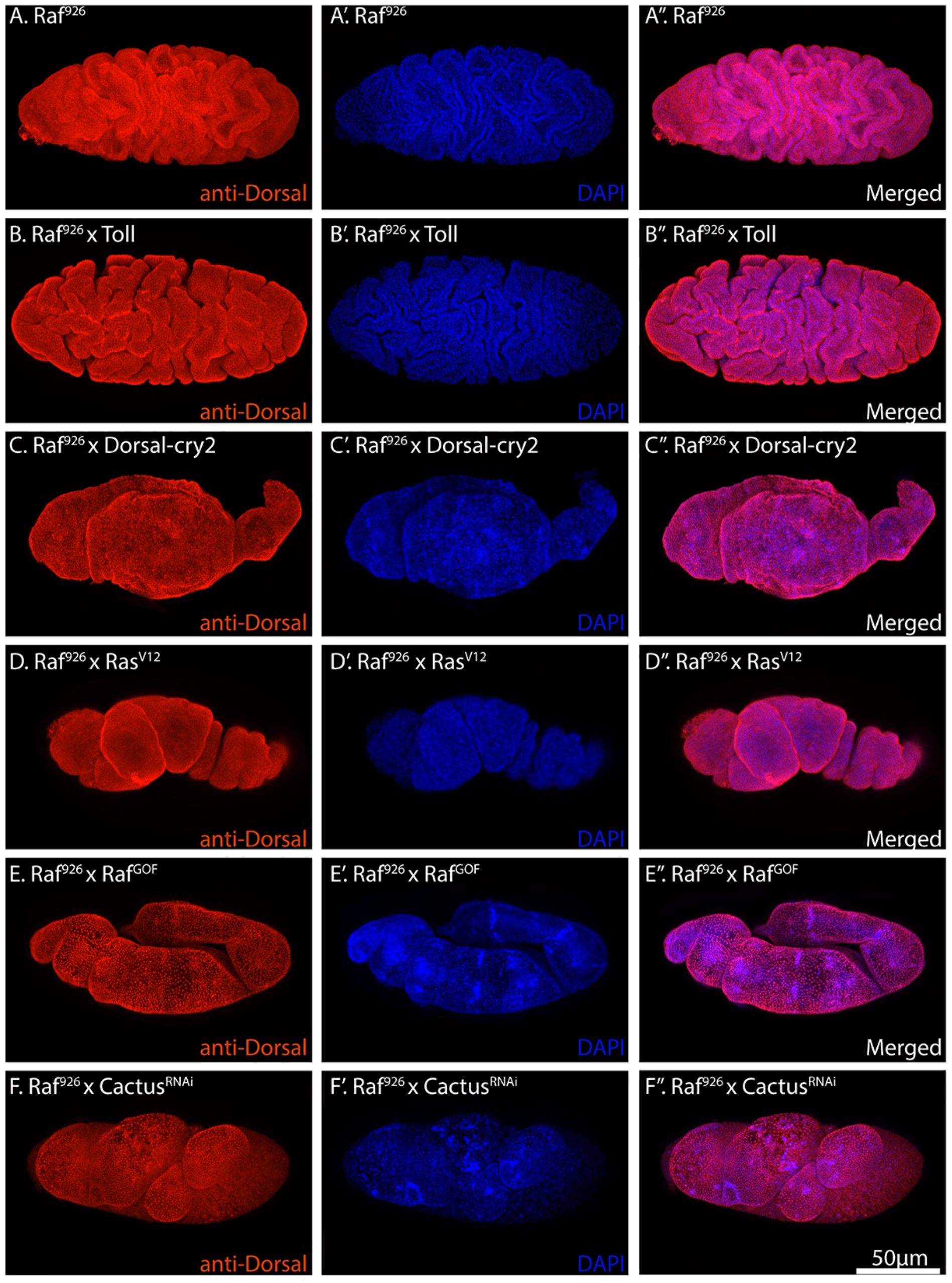
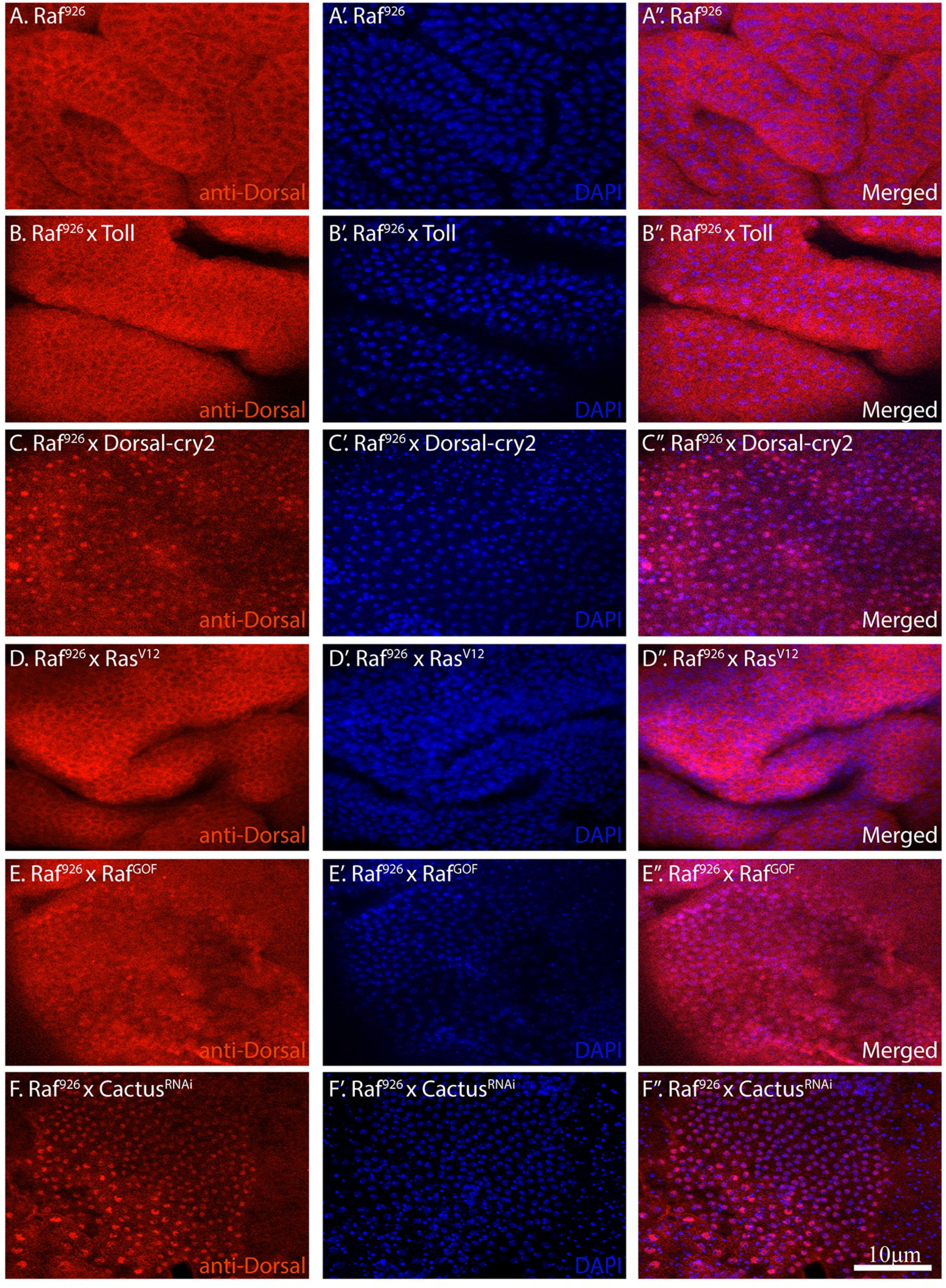
(1 to 5). Analysis of Dorsal nuclear localization in *Raf^926^* mutants. Five figures organized in the same fashion with Dorsal staining in **(A),** nuclear stain in **(A’)** and a merged image in (A’’). (A) shows *Raf^926^* embryos, (B) shows *Raf^926^* embryos crossed to UAS-Toll, (C) shows *Raf^926^* embryos crossed to UAS-Dorsal, (D) shows *Raf^926^* embryos crossed to UAS-Ras^V12^, (E) shows *Raf^926^* embryos crossed to UAS- Raf^GOF^ and (F) shows *Raf^926^* embryos expressing RNAi against *cactus*. (9-1) shows cross sections of early embryos, (9-2) shows surface projections of early embryos, (9-3) shows close-ups of early embryo cross sections, (9-4) shows surface projections of older embryos and (9-5) shows close-ups for late embryos. The figures show little to no Dorsal in the nuclei of early *Raf^926^* embryos except when crossed to Cactus^RNAi^ or Raf^GOF^. This pattern can most readily be seen in older embryos. Overexpressed Dorsal can also be seen in the nuclei of older embryos.

### Resource availability

Further information and requests for resources and reagents should be directed to and will be fulfilled by the Lead Contact Nicholas S. Tolwinski.

### Materials availability

All unique reagents generated in this study are available without any restriction.

## Data availability

RNA-seq data from this study has been deposited to GEO (GSE178187).

## Code availability

## Author contributions

JBL, PK, ICHS, WKL, VYML and NST performed all genetic, cell culture and biochemical experiments. EHZC and NH performed the RNA-seq analysis. JBL and NST wrote the manuscript with input from everyone.

## Acknowledgements

We thank Hugo Bellen and Shinya Yamamoto for allowing us to screen their library and Alison Spencer, Dene Farrell, and Jennifer Zallen for sharing the screen. We thank Benny Shilo for Dorsal-GFP. The antibodies used were obtained from the Developmental Studies Hybridoma Bank, created by the NICHD of the NIH and maintained at The University of Iowa, Department of Biology, Iowa City, IA 52242. Stocks obtained from the Bloomington Drosophila Stock Center (NIH P40OD018537) were used in this study. This work was funded by the Ministry of Education Singapore (Grants: IG18-LR001 and IG19-SI102). EHZC and NH are funded by a start-up grant from Yale-NUS and Duke-NUS.

---

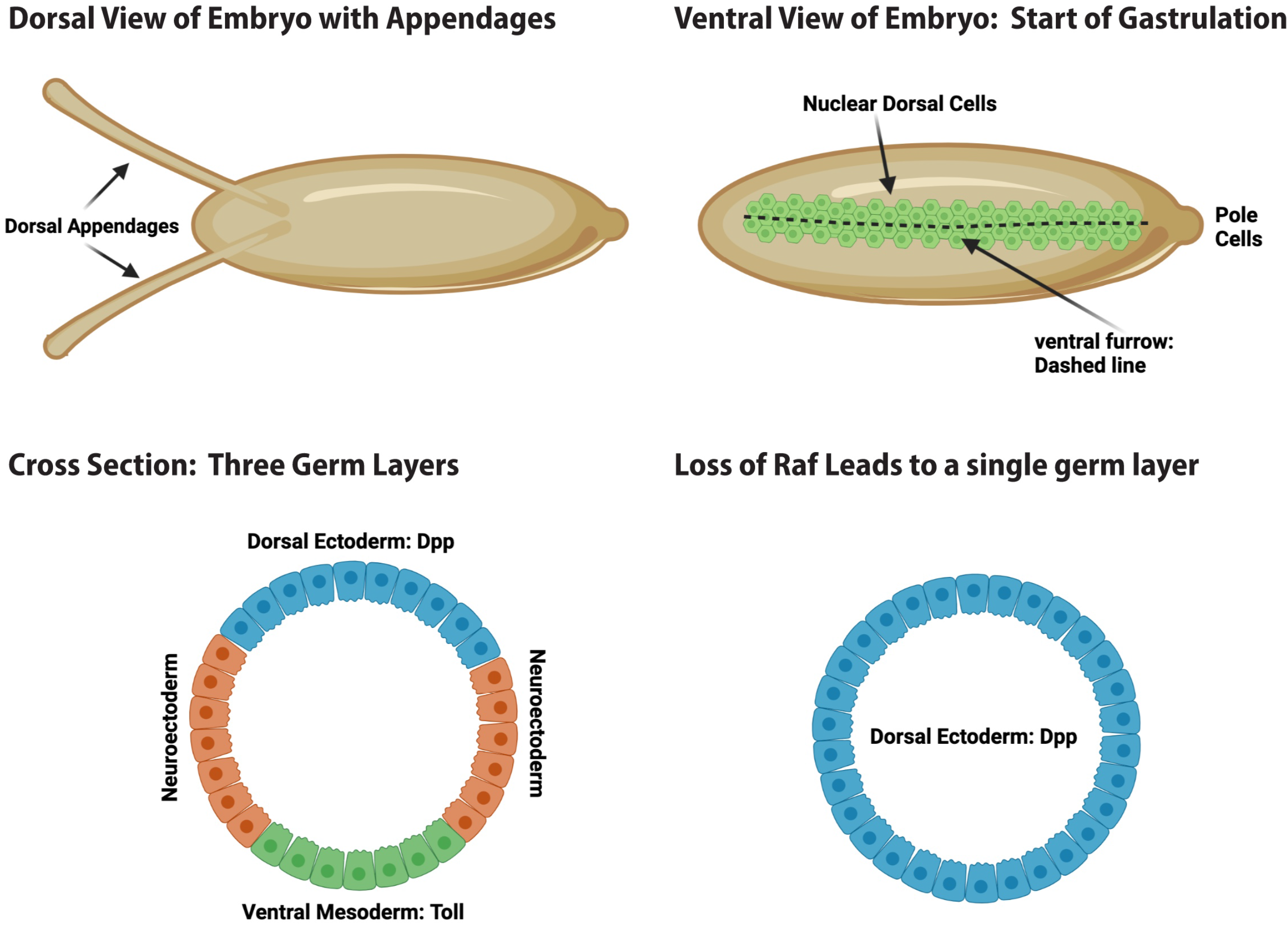

## References

1. Francois V, Solloway M, O’Neill JW, Emery J, Bier E. Dorsal-ventral patterning of the Drosophila embryo depends on a putative negative growth factor encoded by the short gastrulation gene. Genes Dev. 1994;8(21):2602–16. Epub 1994/11/01. PubMed PMID: 7958919.

2. Anderson KV, Bokla L, Nusslein-Volhard C. Establishment of dorsal-ventral polarity in the Drosophila embryo: the induction of polarity by the Toll gene product. Cell. 1985;42(3):791–8. Epub 1985/10/01. PubMed PMID: 3931919.

3. Anderson KV, Jurgens G, Nusslein-Volhard C. Establishment of dorsal-ventral polarity in the Drosophila embryo: genetic studies on the role of the Toll gene product. Cell. 1985;42(3):779–89. Epub 1985/10/01. PubMed PMID: 3931918.

4. Von Ohlen T, Doe CQ. Convergence of dorsal, dpp, and egfr signaling pathways subdivides the drosophila neuroectoderm into three dorsal-ventral columns. Developmental biology. 2000;224(2):362–72.

5. Padgett RW, St Johnston RD, Gelbart WM. A transcript from a Drosophila pattern gene predicts a protein homologous to the transforming growth factor-beta family. Nature. 1987;325(6099):81–4. Epub 1987/01/01. doi: 10.1038/325081a0. PubMed PMID: 3467201.

6. St Johnston RD, Gelbart WM. Decapentaplegic transcripts are localized along the dorsal-ventral axis of the Drosophila embryo. EMBO J. 1987;6(9):2785–91. Epub 1987/09/01. PubMed PMID: 3119329; PubMed Central PMCID: PMCPMC553704.

7. Ferguson EL, Anderson KV. Decapentaplegic acts as a morphogen to organize dorsal-ventral pattern in the Drosophila embryo. Cell. 1992;71(3):451–61. Epub 1992/10/30. PubMed PMID: 1423606.

8. Gabay L, Seger R, Shilo BZ. In situ activation pattern of Drosophila EGF receptor pathway during development. Science. 1997;277(5329):1103–6. Epub 1997/08/22. PubMed PMID: 9262480.

9. Mark GE, MacIntyre RJ, Digan ME, Ambrosio L, Perrimon N. Drosophila melanogaster homologs of the raf oncogene. Mol Cell Biol. 1987;7(6):2134–40. Epub 1987/06/01. PubMed PMID: 3037346; PubMed Central PMCID: PMCPMC365335.

10. Jürgens G, Wieschaus E, Nüsslein-Volhard C, Kluding H. Mutations affecting the pattern of the larval cuticle inDrosophila melanogaster. Wilhelm Roux’s archives of developmental biology. 1984;193(5):283–95. doi: 10.1007/BF00848157.

11. Schejter ED, Shilo B-Z. The Drosophila EGF receptor homolog (DER) gene is allelic to faint little ball, a locus essential for embryonic development. Cell. 1989;56(6):1093–104. doi: https://doi.org/10.1016/0092-8674(89)90642-9.

12. Tearle R, Nusslein-Volhard C. Tubingen mutants and stock list. Dros Inf Serv. 1987;66:209–69.

13. Lindsley DL, Zimm GG. GENES. In: Lindsley DL, Zimm GG, editors. The Genome of Drosophila Melanogaster. San Diego: Academic Press; 1992. p. 1–803.

14. Baek K-H, Fabian JR, Sprenger F, Morrison DK, Ambrosio L. The activity of D-raf in torso signal transduction is altered by serine substitution, N-terminal deletion, and membrane targeting. Developmental biology. 1996;175(2):191–204.

15. Radke K, Johnson K, Guo R, Davidson A, Ambrosio L. Drosophila-raf acts to elaborate dorsoventral pattern in the ectoderm of developing embryos. Genetics. 2001;159(3):1031–44.

16. Duffy JB, Perrimon N. The torso pathway in Drosophila: lessons on receptor tyrosine kinase signaling and pattern formation. Developmental biology. 1994;166(2):380–95.

17. Ambrosio L, Mahowald AP, Perrimon N. l(1)pole hole is required maternally for pattern formation in the terminal regions of the embryo. Development. 1989;106(1):145.

18. Melnick MB, Perkins LA, Lee M, Ambrosio L, Perrimon N. Developmental and molecular characterization of mutations in the Drosophila-raf serine/threonine protein kinase. Development. 1993;118(1):127.

19. Nusslein-Volhard C, Frohnhofer HG, Lehmann R. Determination of anteroposterior polarity in Drosophila. Science. 1987;238(4834):1675-81.

20. Golembo M, Raz E, Shilo B-Z. The Drosophila embryonic midline is the site of Spitz processing, and induces activation of the EGF receptor in the ventral ectoderm. Development. 1996;122(11):3363–70.

21. Golembo M, Schweitzer R, Freeman M, Shilo B-Z. Argos transcription is induced by the Drosophila EGF receptor pathway to form an inhibitory feedback loop. Development. 1996;122(1):223–30.

22. Golembo M, Yarnitzky T, Volk T, Shilo B-Z. Vein expression is induced by the EGF receptor pathway to provide a positive feedback loop in patterning the Drosophila embryonic ventral ectoderm. Genes & development. 1999;13(2):158–62.

23. Banerjee U, Renfranz PJ, Pollock JA, Benzer S. Molecular characterization and expression of sevenless, a gene involved in neuronal pattern formation in the Drosophila eye. Cell. 1987;49(2):281–91.

24. Dickson B, Sprenger F, Morrison D, Hafen E. Raf functions downstream of Rasl in the Sevenless signal transduction pathway. Nature. 1992;360(6404):600-3.

25. Hafen E, Basler K, Edstroem J-E, Rubin GM. Sevenless, a cell-specific homeotic gene of Drosophila, encodes a putative transmembrane receptor with a tyrosine kinase domain. Science. 1987;236(4797):55-63.

26. Perrimon N, Perkins LA. There Must Be 50 Ways to Rule the Signal: The Case of the Drosophila EGF Receptor. Cell. 1997;89(1):13–6. doi: 10.1016/S0092-8674(00)80177-4.

27. Lusk JB, Lam VY, Tolwinski NS. Epidermal Growth Factor Pathway Signaling in Drosophila Embryogenesis: Tools for Understanding Cancer. Cancers (Basel). 2017;9(2). Epub 2017/02/09. doi: 10.3390/cancers9020016. PubMed PMID: 28178204; PubMed Central PMCID: PMCPMC5332939.

28. Sun G, Irvine KD. Chapter Four - Control of Growth During Regeneration. In: Galliot B, editor. Current Topics in Developmental Biology. 108: Academic Press; 2014. p. 95–120.

29. Goff DJ, Nilson LA, Morisato D. Establishment of dorsal-ventral polarity of the Drosophila egg requires capicua action in ovarian follicle cells. Development. 2001;128(22):4553–62.

30. Schüpbach T, Roth S. Dorsoventral patterning in Drosophila oogenesis. Current opinion in genetics & development. 1994;4(4):502–7.

31. Roth S. The origin of dorsoventral polarity in Drosophila. Philosophical Transactions of the Royal Society of London Series B: Biological Sciences. 2003;358(1436):1317–29.

32. Schüpbach T. Germ line and soma cooperate during oogenesis to establish the dorsoventral pattern of egg shell and embryo in Drosophila melanogaster. Cell. 1987;49(5):699–707.

33. Van Buskirk C, Schüpbach T. Versatility in signalling: multiple responses to EGF receptor activation during Drosophila oogenesis. Trends in cell biology. 1999;9(1):1–4.

34. Ruohola-Baker H, Grell E, Chou T-B, Baker D, Jan LY, Jan YN. Spatially localized rhomboid is required for establishment of the dorsal-ventral axis in Drosophila oogenesis. Cell. 1993;73(5):953–65.

35. Morimoto AM, Jordan KC, Tietze K, Britton JS, O’Neill EM, Ruohola-Baker H. Pointed, an ETS domain transcription factor, negatively regulates the EGF receptor pathway in Drosophila oogenesis. Development. 1996;122(12):3745–54.

36. Wasserman JD, Freeman M. An autoregulatory cascade of EGF receptor signaling patterns the Drosophila egg. Cell. 1998;95(3):355–64.

37. Technau M, Knispel M, Roth S. Molecular mechanisms of EGF signaling-dependent regulation of pipe, a gene crucial for dorsoventral axis formation in Drosophila. Development genes and evolution. 2012;222(1):1–17.

38. Belvin MP, Anderson KV. A conserved signaling pathway: the Drosophila toll-dorsal pathway. Annual review of cell and developmental biology. 1996;12(1):393–416.

39. Morisalo D, Anderson KV. Signaling pathways that establish the dorsal-ventral pattern of the Drosophila embryo. Annual review of genetics. 1995;29(1):371–99.

40. Brand AH, Perrimon N. Raf acts downstream of the EGF receptor to determine dorsoventral polarity during Drosophila oogenesis. Genes Dev. 1994;8(5):629–39. Epub 1994/03/01. PubMed PMID: 7926754.

41. Charatsi I, Luschnig S, Bartoszewski S, Nüsslein-Volhard C, Moussian B. Krapfen/dMyd88 is required for the establishment of dorsoventral pattern in the Drosophila embryo. Mechanisms of Development. 2003;120(2):219–26. doi: https://doi.org/10.1016/S0925-4773(02)00410-0.

42. Kambris Z, Bilak H, D’Alessandro R, Belvin M, Imler J-L, Capovilla M. DmMyD88 controls dorsoventral patterning of the Drosophila embryo. EMBO reports. 2003;4(1):64–9. doi: 10.1038/sj.embor.embor714.

43. Horng T, Medzhitov R. Drosophila MyD88 is an adapter in the Toll signaling pathway. Proceedings of the National Academy of Sciences. 2001;98(22):12654–8.

44. Shen B, Manley JL. Pelle kinase is activated by autophosphorylation during Toll signaling in Drosophila. Development. 2002;129(8):1925–33.

45. Towb P, Bergmann A, Wasserman SA. The protein kinase Pelle mediates feedback regulation in the Drosophila Toll signaling pathway. Development. 2001;128(23):4729.

46. Klement J, Rice N, Car B, Abbondanzo S, Powers G, Bhatt P, et al. IkappaBalpha deficiency results in a sustained NF-kappaB response and severe widespread dermatitis in mice. Molecular and cellular biology. 1996;16(5):2341–9.

47. Brown K, Gerstberger S, Carlson L, Franzoso G, Siebenlist U. Control of I kappa B-alpha proteolysis by site-specific, signal-induced phosphorylation. Science. 1995;267(5203):1485-8.

48. Israël A. The IKK complex, a central regulator of NF-κB activation. Cold Spring Harbor perspectives in biology. 2010;2(3):a000158.

49. Park K-J, Krishnan V, O’Malley BW, Yamamoto Y, Gaynor RB. Formation of an IKKα- dependent transcription complex is required for estrogen receptor-mediated gene activation. Molecular cell. 2005;18(1):71–82.

50. Liu J, Sudom A, Min X, Cao Z, Gao X, Ayres M, et al. Structure of the nuclear factor κB- inducing kinase (NIK) kinase domain reveals a constitutively active conformation. Journal of Biological Chemistry. 2012;287(33):27326–34.

51. Hoffmann JA. The immune response of Drosophila. Nature. 2003;426(6962):33.

52. Daigneault J, Klemetsaune L, Wasserman SA. The IRAK Homolog Pelle Is the Functional Counterpart of IκB Kinase in the Drosophila Toll Pathway. PLOS ONE. 2013;8(9):e75150. doi: 10.1371/journal.pone.0075150.

53. Carroll SB, Winslow GM, Twombly VJ, Scott MP. Genes that control dorsoventral polarity affect gene expression along the anteroposterior axis of the Drosophila embryo. Development. 1987;99(3):327–32.

54. Großhans J, Bergmann A, Haffter P, Nüsslein-Volhard C. Activation of the kinase Pelle by Tube in the dorsoventral signal transduction pathway of Drosophila embryo. Nature. 1994;372(6506):563–6.

55. Shelton CA, Wasserman SA. pelle encodes a protein kinase required to establish dorsoventral polarity in the Drosophila embryo. Cell. 1993;72(4):515–25.

56. Sen R, Baltimore D. Inducibility of κ immunoglobulin enhancer-binding protein NF-κB by a posttranslational mechanism. Cell. 1986;47(6):921–8.

57. Nüsslein-Volhard C. Maternal effect mutations that alter the spatial coordinates of the embryo of Drosophila melanogaster. Determinants of spatial organization. 1979;28.

58. Roth S, Stein D, Nüsslein-Volhard C. A gradient of nuclear localization of the dorsal protein determines dorsoventral pattern in the Drosophila embryo. Cell. 1989;59(6):1189–202.

59. Hindley A, Kolch W. Extracellular signal regulated kinase (ERK)/mitogen activated protein kinase (MAPK)-independent functions of Raf kinases. Journal of cell science. 2002;115(8):1575–81.

60. Hipfner DR, Cohen SM. The Drosophila Sterile-20 Kinase Slik Controls Cell Proliferation and Apoptosis during Imaginal Disc Development. PLOS Biology. 2003;1(2):e35. doi: 10.1371/journal.pbio.0000035.

61. Hipfner DR, Cohen SM. Connecting proliferation and apoptosis in development and disease. Nature Reviews Molecular Cell Biology. 2004;5(10):805–15.

62. Matallanas D, Birtwistle M, Romano D, Zebisch A, Rauch J, von Kriegsheim A, et al. Raf Family Kinases: Old Dogs Have Learned New Tricks. Genes & Cancer. 2011;2(3):232–60. doi: 10.1177/1947601911407323. PubMed PMID: 21779496.

63. Yamamoto S, Jaiswal M, Charng WL, Gambin T, Karaca E, Mirzaa G, et al. A drosophila genetic resource of mutants to study mechanisms underlying human genetic diseases. Cell. 2014;159(1):200–14. Epub 2014/09/27. doi: 10.1016/j.cell.2014.09.002. PubMed PMID: 25259927; PubMed Central PMCID: PMCPMC4298142.

64. Perrimon N, Engstrom L, Mahowald AP. Zygotic lethals with specific maternal effect phenotypes in Drosophila melanogaster. I. Loci on the X chromosome. Genetics. 1989;121(2):333–52. Epub 1989/02/01. PubMed PMID: 2499512; PubMed Central PMCID: PMCPMC1203621.

65. Nüsslein-Volhard C, Wieschaus E. Mutations affecting segment number and polarity in Drosophila. Nature. 1980;287(5785):795.

66. Chou T-b, Perrimon N. The autosomal FLP-DFS technique for generating germline mosaics in Drosophila melanogaster. Genetics. 1996;144(4):1673–9.

67. Baker KE, Parker R. Nonsense-mediated mRNA decay: terminating erroneous gene expression. Current Opinion in Cell Biology. 2004;16(3):293–9. doi: https://doi.org/10.1016/j.ceb.2004.03.003.

68. Haelterman NA, Jiang L, Li Y, Bayat V, Sandoval H, Ugur B, et al. Large-scale identification of chemically induced mutations in Drosophila melanogaster. Genome Research. 2014;24(10):1707–18. doi: 10.1101/gr.174615.114.

69. Zirin J, Hu Y, Liu L, Yang-Zhou D, Colbeth R, Yan D, et al. Large-Scale Transgenic Drosophila Resource Collections for Loss- and Gain-of-Function Studies. Genetics. 2020;214(4):755–67. doi: 10.1534/genetics.119.302964.

70. Schneider DS, Hudson KL, Lin TY, Anderson KV. Dominant and recessive mutations define functional domains of Toll, a transmembrane protein required for dorsal-ventral polarity in the Drosophila embryo. Genes Dev. 1991;5(5):797–807. Epub 1991/05/01. PubMed PMID: 1827421.

71. Queenan AM, Ghabrial A, Schupbach T. Ectopic activation of torpedo/Egfr, a Drosophila receptor tyrosine kinase, dorsalizes both the eggshell and the embryo. Development. 1997;124(19):3871–80.

72. Huisken J, Swoger J, Del Bene F, Wittbrodt J, Stelzer EH. Optical sectioning deep inside live embryos by selective plane illumination microscopy. Science. 2004;305(5686):1007-9.

73. Voie AH, Burns D, Spelman F. Orthogonal!plane fluorescence optical sectioning: Three! dimensional imaging of macroscopic biological specimens. Journal of microscopy. 1993;170(3):229–36.

74. Sweeton D, Parks S, Costa M, Wieschaus E. Gastrulation in Drosophila: the formation of the ventral furrow and posterior midgut invaginations. Development. 1991;112(3):775–89.

75. Hartenstein V, Lee A, Toga AW. A graphic digital database of Drosophila embryogenesis. Trends in genetics. 1995;11(2):51–8.

76. Ganguly A, Ip YT. Change of Epithelial Fate. Rise and Fall of Epithelial Phenotype: Springer; 2005. p. 101–10.

77. Schüpbach T, Wieschaus E. Maternal-effect mutations altering the anterior-posterior pattern of the Drosophila embryo. Roux’s archives of developmental biology. 1986;195(5):302–17.

78. Ambrosio L, Mahowald AP, Perrimon N. Requirement of the Drosophila raf homologue for torso function. Nature. 1989;342(6247):288–91.

79. Irvine KD, Wieschaus E. Cell intercalation during Drosophila germband extension and its regulation by pair-rule segmentation genes. Development. 1994;120(4):827–41.

80. Gabay L, Seger R, Shilo BZ. MAP kinase in situ activation atlas during Drosophila embryogenesis. Development. 1997;124(18):3535–41.

81. Stathopoulos A, Van Drenth M, Erives A, Markstein M, Levine M. Whole-Genome Analysis of Dorsal-Ventral Patterning in the *Drosophila* Embryo. Cell. 2002;111(5):687–701. doi: 10.1016/S0092-8674(02)01087-5.

82. Koenecke N, Johnston J, Gaertner B, Natarajan M, Zeitlinger J. Genome-wide identification of Drosophila dorso-ventral enhancers by differential histone acetylation analysis. Genome Biology. 2016;17(1):196. doi: 10.1186/s13059-016-1057-2.

83. Koenecke N, Johnston J, He Q, Meier S, Zeitlinger J. Drosophila poised enhancers are generated during tissue patterning with the help of repression. Genome research. 2017;27(1):64–74.

84. DeLotto R, DeLotto Y, Steward R, Lippincott-Schwartz J. Nucleocytoplasmic shuttling mediates the dynamic maintenance of nuclear Dorsal levels during Drosophila embryogenesis. Development. 2007;134(23):4233–41.

85. Hong J-W, Hendrix DA, Papatsenko D, Levine MS. How the Dorsal gradient works: insights from postgenome technologies. Proceedings of the National Academy of Sciences. 2008;105(51):20072–6.

86. Ip YT, Park RE, Kosman D, Yazdanbakhsh K, Levine M. dorsal-twist interactions establish snail expression in the presumptive mesoderm of the Drosophila embryo. Genes & development. 1992;6(8):1518–30.

87. Shirokawa JM, Courey AJ. A direct contact between the dorsal rel homology domain and Twist may mediate transcriptional synergy. Molecular and cellular biology. 1997;17(6):3345–55.

88. Thisse B, Messal ME, Perrin-Schmitt F. The twist gene: isolation of a Drosophila zygotle gene necessary for the establishment of dorsoventral pattern. Nucleic acids research. 1987;15(8):3439–53.

89. Sandmann T, Girardot C, Brehme M, Tongprasit W, Stolc V, Furlong EE. A core transcriptional network for early mesoderm development in Drosophila melanogaster. Genes & development. 2007;21(4):436–49.

90. Bate M, Rushton E, Currie DA. Cells with persistent twist expression are the embryonic precursors of adult muscles in Drosophila. Development. 1991;113(1):79–89.

91. Li S, Sedivy JM. Raf-1 protein kinase activates the NF-kappa B transcription factor by dissociating the cytoplasmic NF-kappa BI kappa B complex. Proceedings of the National Academy of Sciences. 1993;90(20):9247–51.

92. Baldi L, Brown K, Franzoso G, Siebenlist U. Critical role for lysines 21 and 22 in signal-induced, ubiquitin-mediated proteolysis of I kappa B-alpha. J Biol Chem. 1996;271(1):376–9. Epub 1996/01/05. PubMed PMID: 8550590.

93. Johnson HE, Goyal Y, Pannucci NL, Schüpbach T, Shvartsman SY, Toettcher JE. The Spatiotemporal Limits of Developmental Erk Signaling. Developmental Cell. 2017;40(2):185–92. doi: https://doi.org/10.1016/j.devcel.2016.12.002.

94. Johnson HE, Toettcher JE. Signaling dynamics control cell fate in the early Drosophila embryo. Developmental cell. 2019;48(3):361–70. e3.

95. Keenan SE, Blythe SA, Marmion RA, Djabrayan NJ-V, Wieschaus EF, Shvartsman SY. Rapid Dynamics of Signal-Dependent Transcriptional Repression by Capicua. Developmental Cell. 2020.

96. McFann S, Dutta S, Toettcher JE, Shvartsman SY. Temporal integration of inductive cues on the way to gastrulation. Proceedings of the National Academy of Sciences. 2021;118(23):e2102691118. doi: 10.1073/pnas.2102691118.

97. Helman A, Lim B, Andreu MJ, Kim Y, Shestkin T, Lu H, et al. RTK signaling modulates the Dorsal gradient. Development. 2012;139(16):3032–9. doi: 10.1242/dev.075812.

98. Papagianni A, Forés M, Shao W, He S, Koenecke N, Andreu MJ, et al. Capicua controls Toll/IL-1 signaling targets independently of RTK regulation. Proceedings of the National Academy of Sciences. 2018;115(8):1807–12. doi: 10.1073/pnas.1713930115.

99. Li S, Sedivy JM. Raf-1 protein kinase activates the NF-kappa B transcription factor by dissociating the cytoplasmic NF-kappa B-I kappa B complex. Proceedings of the National Academy of Sciences. 1993;90(20):9247. doi: 10.1073/pnas.90.20.9247.

100. Zarnegar B, Yamazaki S, He JQ, Cheng G. Control of canonical NF-κB activation through the NIK–IKK complex pathway. Proceedings of the National Academy of Sciences. 2008;105(9):3503. doi: 10.1073/pnas.0707959105.

101. Lynch JA, Peel AD, Drechsler A, Averof M, Roth S. EGF Signaling and the Origin of Axial Polarity among the Insects. Current Biology. 2010;20(11):1042–7. doi: 10.1016/j.cub.2010.04.023.

102. St Johnston D, Nüsslein-Volhard C. The origin of pattern and polarity in the Drosophila embryo. Cell. 1992;68(2):201–19.

103. Brennan CA, Anderson KV. Drosophila: The Genetics of Innate Immune Recognition and Response. Annual Review of Immunology. 2004;22(1):457–83. doi: 10.1146/annurev.immunol.22.012703.104626. PubMed PMID: 15032585.

104. Charles A. Janeway J, Medzhitov R. Innate Immune Recognition. Annual Review of Immunology. 2002;20(1):197–216. doi: 10.1146/annurev.immunol.20.083001.084359. PubMed PMID: 11861602.

105. Kawai T, Akira S. The role of pattern-recognition receptors in innate immunity: update on Toll-like receptors. Nature Immunology. 2010;11:373. doi: 10.1038/ni.1863.

106. Leulier F, Lemaitre B. Toll-like receptors — taking an evolutionary approach. Nature Reviews Genetics. 2008;9:165. doi: 10.1038/nrg2303.

107. Xu T, Rubin GM. Analysis of genetic mosaics in developing and adult Drosophila tissues. Development. 1993;117(4):1223–37.

108. Lee T, Feig L, Montell DJ. Two distinct roles for Ras in a developmentally regulated cell migration. Development. 1996;122(2):409–18. Epub 1996/02/01. PubMed PMID: 8625792.

109. Maxton-Küchenmeister J, Handel K, Schmidt-Ott U, Roth S, Jäckle H. Toll homolog expression in the beetle Tribolium suggests a different mode of dorsoventral patterning than in Drosophila embryos. Mechanisms of Development. 1999;83(1):107–14. doi: https://doi.org/10.1016/S0925-4773(99)00041-6.

110. Perkins LA, Holderbaum L, Tao R, Hu Y, Sopko R, McCall K, et al. The Transgenic RNAi Project at Harvard Medical School: Resources and Validation. Genetics. 2015;201(3):843. doi: 10.1534/genetics.115.180208.

111. Halfon MS, Gisselbrecht S, Lu J, Estrada B, Keshishian H, Michelson AM. New fluorescent protein reporters for use with the drosophila gal4 expression system and for vital detection of balancer chromosomes. genesis. 2002;34(112):135–8. doi: 10.1002/gene.10136.

112. Riabinina O, Luginbuhl D, Marr E, Liu S, Wu MN, Luo L, et al. Improved and expanded Q-system reagents for genetic manipulations. Nature methods. 2015;12(3):219.

113. Bunnag N, Tan QH, Kaur P, Ramamoorthy A, Sung ICH, Lusk J, et al. An Optogenetic Method to Study Signal Transduction in Intestinal Stem Cell Homeostasis. J Mol Biol. 2020. Epub 2020/03/24. doi: 10.1016/j.jmb.2020.03.019. PubMed PMID: 32201167.

114. Jankovics F, Henn L, Bujna Á, Vilmos P, Kiss N, Erdélyi M. A Functional Genomic Screen Combined with Time-Lapse Microscopy Uncovers a Novel Set of Genes Involved in Dorsal Closure of Drosophila Embryos. PLOS ONE. 2011;6(7):e22229. doi: 10.1371/journal.pone.0022229.

115. Potter CJ, Luo L. Using the Q system in Drosophila melanogaster. Nature Protocols. 2011;6:1105. doi: 10.1038/nprot.2011.347 https://www.nature.com/articles/nprot.2011.347#supplementary-information.

116. Venken KJT, Popodi E, Holtzman SL, Schulze KL, Park S, Carlson JW, et al. A Molecularly Defined Duplication Set for the X Chromosome of *Drosophila melanogaster*. Genetics. 2010;186(4):1111. doi: 10.1534/genetics.110.121285.

117. Bischof J, Bjorklund M, Furger E, Schertel C, Taipale J, Basler K. A versatile platform for creating a comprehensive UAS-ORFeome library in Drosophila. Development. 2013;140(11):2434–42. doi: 10.1242/dev.088757. PubMed PMID: 23637332.

118. Richens JH, Barros TP, Lucas EP, Peel N, Pinto DMS, Wainman A, et al. The Drosophila Pericentrin-like-protein (PLP) cooperates with Cnn to maintain the integrity of the outer PCM. Biol Open. 2015;4(8):1052–61. doi: 10.1242/bio.012914. PubMed PMID: 26157019.

119. Groth AC, Fish M, Nusse R, Calos MP. Construction of transgenic Drosophila by using the site-specific integrase from phage phiC31. Genetics. 2004;166(4):1775–82. PubMed PMID: 15126397; PubMed Central PMCID: PMCPMC1470814.

120. Kaur P, Saunders TE, Tolwinski NS. Coupling optogenetics and light-sheet microscopy, a method to study Wnt signaling during embryogenesis. Sci Rep. 2017;7(1):16636. Epub 2017/12/02. doi: 10.1038/s41598-017-16879-0. PubMed PMID: 29192250; PubMed Central PMCID: PMCPMC5709371.

121. Muller HA, Wieschaus E. armadillo, bazooka, and stardust are critical for early stages in formation of the zonula adherens and maintenance of the polarized blastoderm epithelium in Drosophila. J Cell Biol. 1996;134(1):149–63. Epub 1996/07/01. PubMed PMID: 8698811.

122. Tolwinski NS, Wieschaus E. Armadillo nuclear import is regulated by cytoplasmic anchor Axin and nuclear anchor dTCF/Pan. Development. 2001;128(11):2107–17. Epub 2001/08/09. PubMed PMID: 11493532.

123. Colosimo PF, Tolwinski NS. Wnt, Hedgehog and junctional Armadillo/beta-catenin establish planar polarity in the Drosophila embryo. PLoS One. 2006;1:e9. Epub 2006/12/22. doi: 10.1371/journal.pone.0000009. PubMed PMID: 17183721; PubMed Central PMCID: PMCPMC1762359.

124. Dobin A, Davis CA, Schlesinger F, Drenkow J, Zaleski C, Jha S, et al. STAR: ultrafast universal RNA-seq aligner. Bioinformatics. 2013;29(1):15–21.

125. Li B, Dewey CN. RSEM: accurate transcript quantification from RNA-Seq data with or without a reference genome. BMC Bioinformatics. 2011;12(1):323. doi: 10.1186/1471-2105-12-323.

126. Love MI, Huber W, Anders S. Moderated estimation of fold change and dispersion for RNA-seq data with DESeq2. Genome Biology. 2014;15(12):550. doi: 10.1186/s13059-014-0550-8.

127. Xie Z, Bailey A, Kuleshov MV, Clarke DJB, Evangelista JE, Jenkins SL, et al. Gene Set Knowledge Discovery with Enrichr. Current Protocols. 2021;1(3):e90. doi: https://doi.org/10.1002/cpz1.90.

128. Kuleshov MV, Jones MR, Rouillard AD, Fernandez NF, Duan Q, Wang Z, et al. Enrichr: a comprehensive gene set enrichment analysis web server 2016 update. Nucleic Acids Res. 2016;44(W1):W90–7. Epub 2016/05/05. doi: 10.1093/nar/gkw377. PubMed PMID: 27141961; PubMed Central PMCID: PMCPMC4987924.

129. Chen EY, Tan CM, Kou Y, Duan Q, Wang Z, Meirelles GV, et al. Enrichr: interactive and collaborative HTML5 gene list enrichment analysis tool. BMC Bioinformatics. 2013;14:128. Epub 2013/04/17. doi: 10.1186/1471-2105-14-128. PubMed PMID: 23586463; PubMed Central PMCID: PMCPMC3637064.

130. Subramanian A, Tamayo P, Mootha VK, Mukherjee S, Ebert BL, Gillette MA, et al. Gene set enrichment analysis: A knowledge-based approach for interpreting genome-wide expression profiles. Proceedings of the National Academy of Sciences. 2005;102(43):15545. doi: 10.1073/pnas.0506580102.

131. Liberzon A, Subramanian A, Pinchback R, Thorvaldsdóttir H, Tamayo P, Mesirov JP. Molecular signatures database (MSigDB) 3.0. Bioinformatics. 2011;27(12):1739–40. doi: 10.1093/bioinformatics/btr260.

132. Liberzon A, Birger C, Thorvaldsdóttir H, Ghandi M, Mesirov JP, Tamayo P. The Molecular Signatures Database (MSigDB) hallmark gene set collection. Cell Syst. 2015;1(6):417–25. doi: 10.1016/j.cels.2015.12.004. PubMed PMID: 26771021.

133. Yu G, Wang L-G, Han Y, He Q-Y. clusterProfiler: an R Package for Comparing Biological Themes Among Gene Clusters. OMICS: A Journal of Integrative Biology. 2012;16(5):284–7. doi: 10.1089/omi.2011.0118.

